# Foreign body responses in central nervous system mimic natural wound responses and alter biomaterial functions

**DOI:** 10.1101/797092

**Authors:** Timothy M. O’Shea, Alexander L. Wollenberg, Jae H. Kim, Yan Ao, Timothy J. Deming, Michael V. Sofroniew

## Abstract

Biomaterials hold promise for diverse therapeutic applications in the central nervous system (CNS). Little is known about molecular factors that determine CNS foreign body responses (FBRs) *in vivo*, or about how such responses influence biomaterial function. Here, we probed these factors using a platform of injectable hydrogels readily modified to present interfaces with different representative physiochemical properties to host cells. We show that biomaterial FBRs mimic specialized multicellular CNS wound responses not present in peripheral tissues, which serve to isolate damaged neural tissue and restore barrier functions. Moreover, we found that the nature and intensity of CNS FBRs are determined by definable properties. For example, cationic, anionic or nonionic interfaces with CNS cells elicit quantifiably different levels of stromal cell infiltration, inflammation, neural damage and amyloid production. The nature and intensity of FBRs significantly influenced hydrogel resorption and molecular delivery functions. These results characterize specific molecular mechanisms that drive FBRs in the CNS and have important implications for developing effective biomaterials for CNS applications.

## Introduction

Biomaterials are under widespread investigation for experimental and therapeutic applications in the central nervous system (CNS) [1]. Biomaterials with specialized properties have been incorporated in implantable neuroprostheses that are used clinically for recording neuronal activity and stimulating neural circuits in the CNS [2, 3]. In addition, injectable biomaterial hydrogels are used extensively as experimental tools to provide local delivery of growth factors for example to attract injured and regenerating axons to grow across CNS lesions [4, 5] as well as to afford molecular and physical support to co-suspended neural progenitor cells (NPC) to improve survival and modulate differentiation upon grafting into the CNS [6–8].

While various classes of biomaterials may be used for applications in the CNS, synthetic or naturally derived polymers are the predominate choice for directly interfacing with host CNS tissue and cells [9, 10]. For neuroprostheses, diverse classes of polymers are used as the outer substrates or coatings for metal electrodes intended to improve device insertion and long term performance *in vivo* [10, 11]. In addition, new devices based on electrically conducting polymers are also being developed [12]. For hydrogels, considerable progress has been made in synthesizing new polymers with tunable physiochemical properties to achieve spatial and temporal control of molecular release or provide unique support to co-suspended cells [13–16].

Despite innovations in the many engineering and design aspects of biomaterials that are used for CNS applications, little is known about factors that determine the CNS foreign body response (FBR) *in vivo* to these materials, or how such responses influence function. Assessments of cytotoxicity and biocompatibility of biomaterial interfaces with host cells, or biomaterial functions such as molecular release kinetics are often evaluated primarily *in vitro* or through subcutaneous implantation *in vivo* [17, 18]. However, *in vitro* observations and the FBR associated with subcutaneously implanted biomaterials need not predict *in vivo* performance in the CNS, where there is a highly specialized and unique multicellular wound response that serves to isolate damaged tissue and restore barrier functions.

Substantial advances have been made in understanding the CNS wound response, which can inform the study of CNS FBRs. The CNS wound response is a biologically conserved process whereby perturbations of CNS tissue lead to dynamic multi-cellular interactions that efficiently isolate regions perceived as damaged, infected or diseased to protect adjacent viable neural tissue and permit recruited inflammation to resolve potentially noxious elements [5, 19–21]. This multicellular CNS wound response generates tissue lesions characterized by distinct non-neural and neural cellular compartments in which non-neural lesion cores with stromal and inflammatory cells are segregated from adjacent spared and viable neural tissue by an astroglial scar or limitans border [22]. The cellular responses to materials perceived as foreign are likely to be related to this multicellular natural wound response [23], but the study of CNS FBRs to biomaterials *in vivo* has not kept pace with recent advances made in the cell biology of CNS injury. There is a need to better understand: (i) key cellular interactions of CNS FBRs to biomaterials, (ii) biomaterial properties that influence the severity of CNS FBRs, and (iii) the extent to which CNS FBRs influence biomaterial function *in vivo*. Here, we evaluated these parameters by comparing multiple synthetic and naturally derived hydrogel-based biomaterials.

As a platform to determine the effects of specific molecular features on biomaterial-evoked FBRs, we used synthetic, diblock copolypeptide hydrogels (DCH), which can be readily modified to present interfaces with different representative physiochemical properties to host cells whilst retaining consistent polymer chain lengths and mechanical properties [8, 24, 25]. These shear-thinning, physical hydrogels can be administered to focal CNS regions in a minimally invasive manner by infusion through narrow cannulae ensuring that minimal tissue damage is induced by the implantation of the biomaterial that could otherwise confound the CNS FBR evaluation of the host-biomaterial interface [24–26]. Moreover, because these hydrogels can be prepared to exhibit the same mechanical stiffness as CNS tissue (ca. 100-400 Pa) [25], they can be fixed and processed *in situ* and retain their intimate contact with immediately adjacent host cells so that directly contiguous biomaterial-host interfaces can be examined and quantified in detail. This type of analysis is not possible with rigid, mechanically stiff or hydrophobic materials and coated electrodes that must be removed before tissue sectioning, because this removal invariably also disrupts the immediate material-cellular interfaces, which cannot then be evaluated.

We benchmarked our findings using DCH that present physiochemically defined interfaces against various commonly used, commercially available and previously CNS tested hydrogel formulations and compared FBRs with naturally evoked CNS wound responses modeled using chemically induced ischemic strokes that were equivalent in size to the hydrogel depots. We show that the CNS FBR to biomaterial interfaces is a multicellular process that mimics conserved elements of CNS wound healing and exists on a definable severity spectrum characterized by the presence or absence of specific different types of non-neural cells. Furthermore, the nature and intensity of CNS FBRs are determined by definable material properties and elicits quantifiable differences in relevant functions including hydrogel resorption and molecular delivery.

## Results

### Hydrogel-evoked FBRs vary and mimic CNS wound responses

We first compared cellular profiles of mild or severe CNS wound responses with FBRs of two structurally similar, synthetic, diblock copolypeptide hydrogels (DCH) that present either strongly cationic (DCH_K_) or nonionic (DCH_MO_) interfaces to host cells (Fig. 1a). Lysine-based DCH_K_ and methionine-sulfoxide based DCH_MO_ are both well tolerated *in vivo* [8, 25], but exhibit noticeably different FBRs. For comparison, mild or severe CNS wound responses were induced by injection of innocuous phosphate buffered saline (PBS), or N5-(1-iminoethyl)-l-ornithine (L-NIO) solution that creates a small focal ischemic stroke [27]. PBS, DCH_MO_, DCH_K_, or L-NIO were injected into caudate putamen (CP) of mouse forebrain (Fig. 1b). The CP site is easily accessible, allows for consistent and reproducible hydrogel injections that are well tolerated by the mice, and is composed of neural tissue with neuronal cell bodies, myelinated axon bundles and a diversity of neuroglial making it an advantageous location for standardized CNS FBR assessments [25]. Hydrogel FBRs and wound responses were characterized with immunohistochemistry for CD13 to identify non-neural cells including stromal and peripheral myeloid lineage cells recruited as part of the sterile inflammatory response to tissue damage [28]. CD13 positively identifies the full temporal range of non-neural tissue remodeling from acute inflammation through to chronic fibrotic lesion formation. In the healthy uninjured mouse brain, CD13 is expressed by pericytes and perivascular fibroblasts along cerebrovasculature and meningeal fibroblasts [29] (Supplementary Fig. 1a-d). CD13 is not expressed by neuroglia or neurons. Staining for glial fibrillary acidic protein (GFAP) identified reactive and scar-border forming astrocytes [30], P2RY12 identified CNS microglia and NeuN identified viable neurons.

**Fig. 1.**
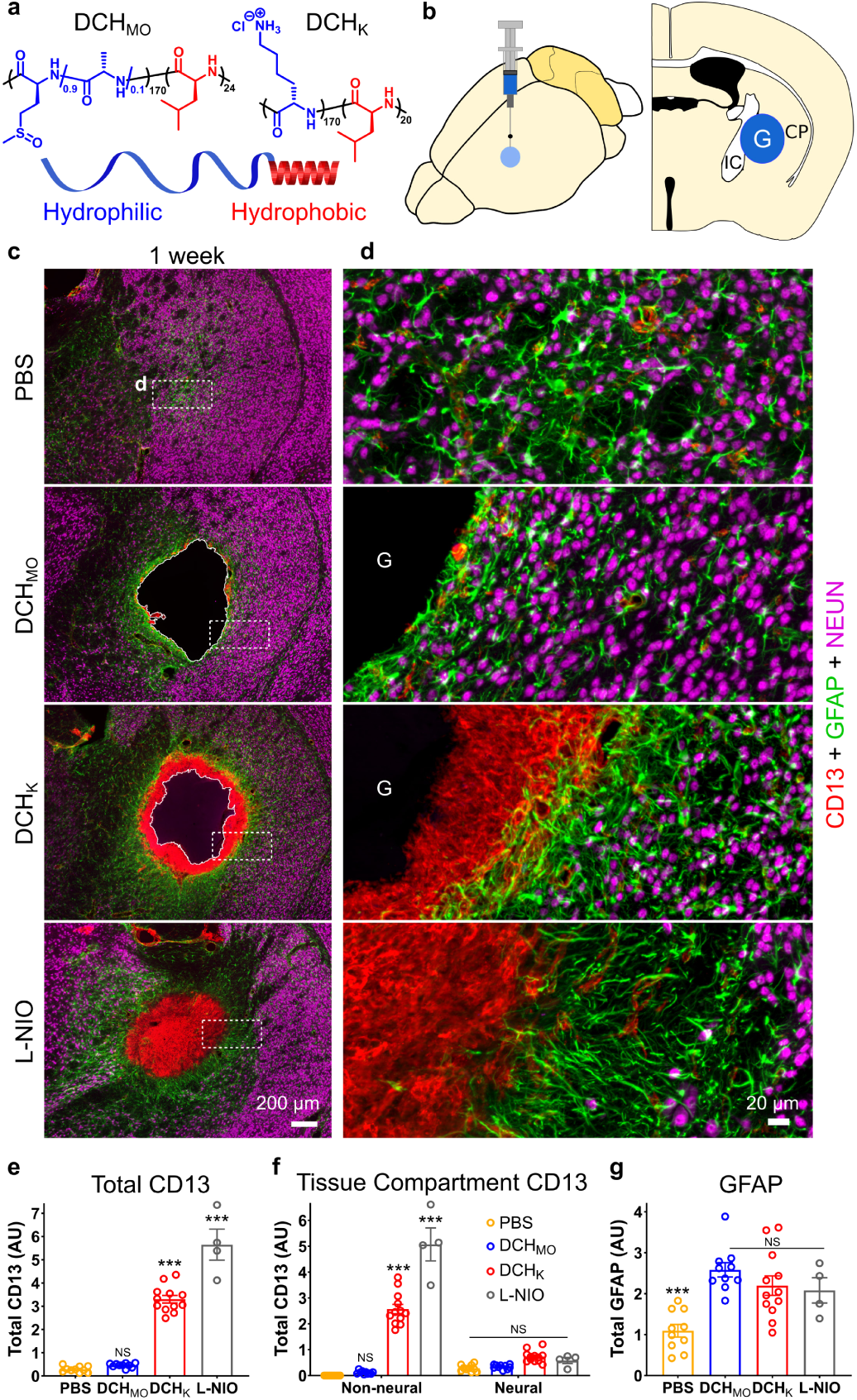
Hydrogel-evoked FBRs vary and exhibit cellular features of CNS wound response. **a**. Schematic and chemical structures of synthetic hydrogels used as tools to study CNS FBR. **b.** Experimental model of *in vivo* injections into caudate putamen (CP) of mouse forebrain. **c,d.** Survey and detail images of stromal and inflammatory cells (CD13), astrocytes (GFAP) and neurons (NeuN) at one week after injections of PBS, hydrogels or L-NIO-induced stroke. **e-g.** Quantification of total immunohistochemical staining. **e.** Total CD13 in CP. NS not significant, ***P < 0.0001 versus PBS injection, one-way ANOVA with Bonferroni. **f.** Total CD13 in either non-neural or neural tissue compartments. NS or ***P < 0.0001 versus PBS injection, two-way ANOVA with Bonferroni. **g.** Total GFAP in CP. NS or ***P < 0.0001 versus DCH_MO_, one-way ANOVA with Bonferroni. Graphs are mean ± s.e.m with individual data points showing *n* mice per group. AU arbitrary units, G hydrogel, IC internal capsule. (one column width figure)

At one week after injections, (i) PBS infusions were detectible as small areas of GFAP-positive reactive astrocytes; (ii) nonionic DCH_MO_ and cationic DCH_K_, exhibited similarly sized deposits; and (iii) L-NIO created sharply demarcated areas of infarcted tissue equivalent in size to hydrogel deposits (Fig. 1c). DCH_MO_ and DCH_K_ deposits and L-NIO infarcts were all surrounded by similarly appearing borders of GFAP-positive astrocytes that clearly separated neural tissue containing NeuN-positive neurons from non-neural cores with CD13-positive cells and hydrogels (Fig. 1c,d). CD13-positive cells were not elevated at the centers of PBS injections or at interfaces between DCH_MO_ and host. By contrast, a dense CD13-positive cell capsule, 164±19µm in thickness, had completely surrounded and infiltrated the margins of DCH_K_ deposits, while CD13-positive cells had entirely filled L-NIO infarcts (Fig. 1c,d). Similar CD13-positive cell accumulation and non-neural/neural cell compartmentalization was observed one week after forebrain stab injury through cerebral cortex (Supplementary Fig.1e). Quantification showed that total CD13 levels across the different compartments of infusion sites and adjacent neural tissue were similar after PBS and DCH_MO_ and were significantly higher after DCH_K_ and L-NIO (Fig. 1e; Supplementary Fig. 2). Notably, the significant increases in CD13 levels after DCH_K_ and L-NIO were confined to central non-neural tissue compartments of hydrogel plus infiltrating cells, and there was no significant difference in CD13 levels in surrounding neural tissue (defined as tissue containing neuroglia and neurons) adjacent to PBS, DCH_MO_, DCH_K_ or L-NIO (Fig. 1f). Moreover, although GFAP levels were significantly lower adjacent to PBS injections, there was no significant difference in GFAP levels adjacent to DCH_MO_, DCH_K_ or L-NIO (Fig. 1g).

**Fig. 2.**
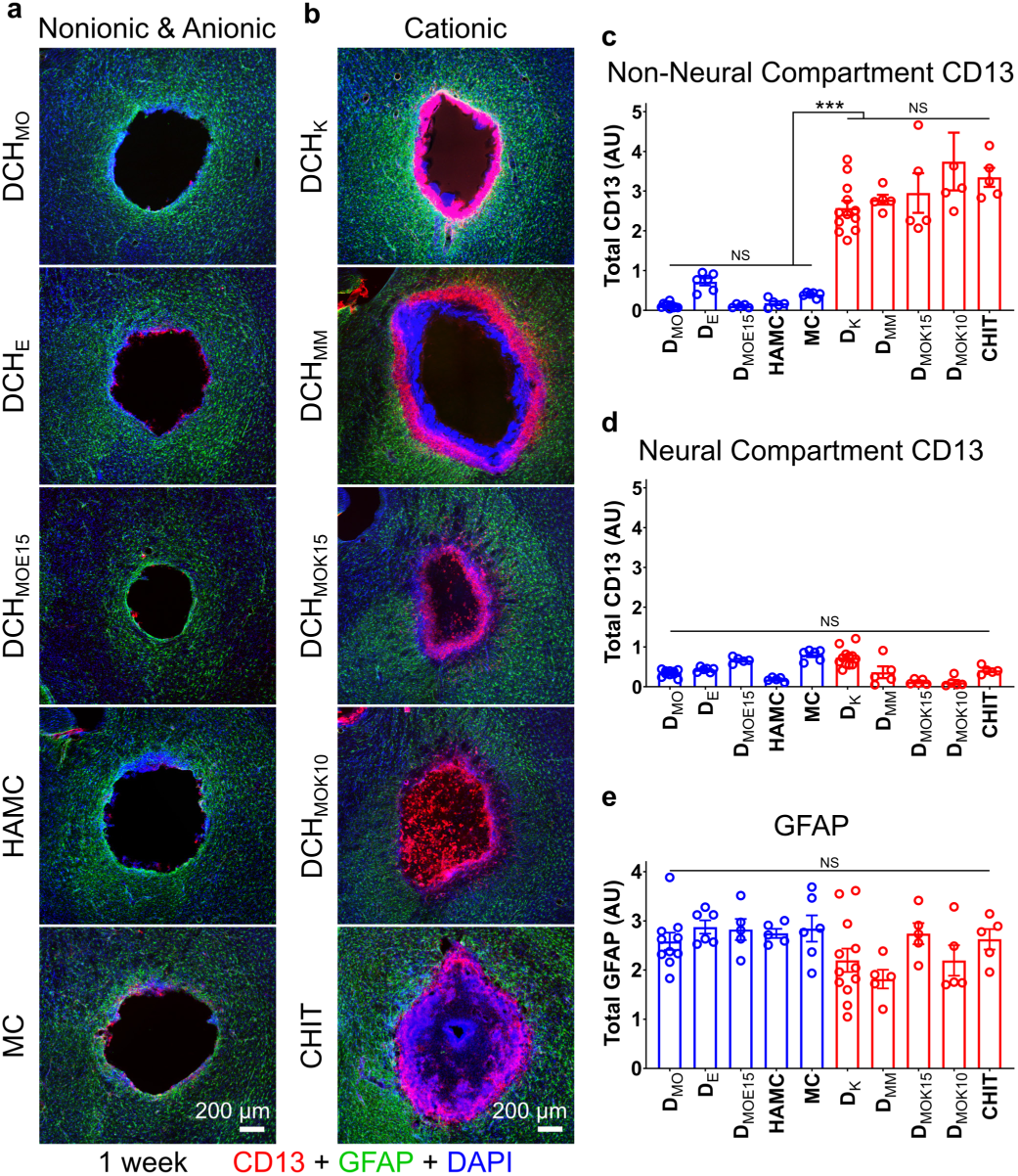
Hydrogels that present cationic interfaces with host cells exhibit increased FBR severity. **a,b.** Survey images of stromal and inflammatory cells (CD13), astrocytes (GFAP) and all cell nuclei (DAPI) at one week after injection of nonionic and anionic (a), and cationic (b) hydrogels. **c-e.** Quantification of total immunohistochemical staining. **c,d.** Total CD13 in non-neural (c) and neural (d) tissue compartments. NS not significant, ***P < 0.0001, two-way ANOVA with Bonferroni. **e.** Total GFAP in CP, NS, one-way ANOVA with Bonferroni. Graphs are mean ± s.e.m with individual data points showing *n* mice per group. (one column width figure)

These findings show that DCH-based hydrogels presenting precisely defined cationic (DCH_K_) or nonionic (DCH_MO_) interfaces to host cells exhibit FBRs whose cellular profiles and compartmentalization mimic those of normal CNS wound responses, and that the FBR to different hydrogels can vary significantly. These findings also suggested that hydrogel surface chemistry and properties such as charge may influence hydrogel FBR.

### FBRs vary with definable hydrogel properties

We next characterized FBRs to various cationic, anionic and nonionic hydrogels. We manufactured DCH that presented different polar side chains to interface with host cells but maintained consistent poly(L-leucine) hydrophobic blocks [8, 24, 25] (Supplementary Fig. 3a,b). We compared these DCH with various commercially available formulations commonly used in CNS applications: methylcellulose (MC) [31], a hyaluronic acid/methylcellulose blend (HAMC) [32], and a chitosan/β-glycerophosphate (CHIT) system [33]. Hydrogels with comparable mechanical properties (see [8, 25, 34] and Supplementary Fig.3c) were injected into mouse forebrain (Fig. 1b), and cellular FBR profiles were first characterized with immunohistochemistry for CD13, GFAP and NeuN (Fig. 2; Supplementary Figs. 2,4,5).

**Fig. 3.**
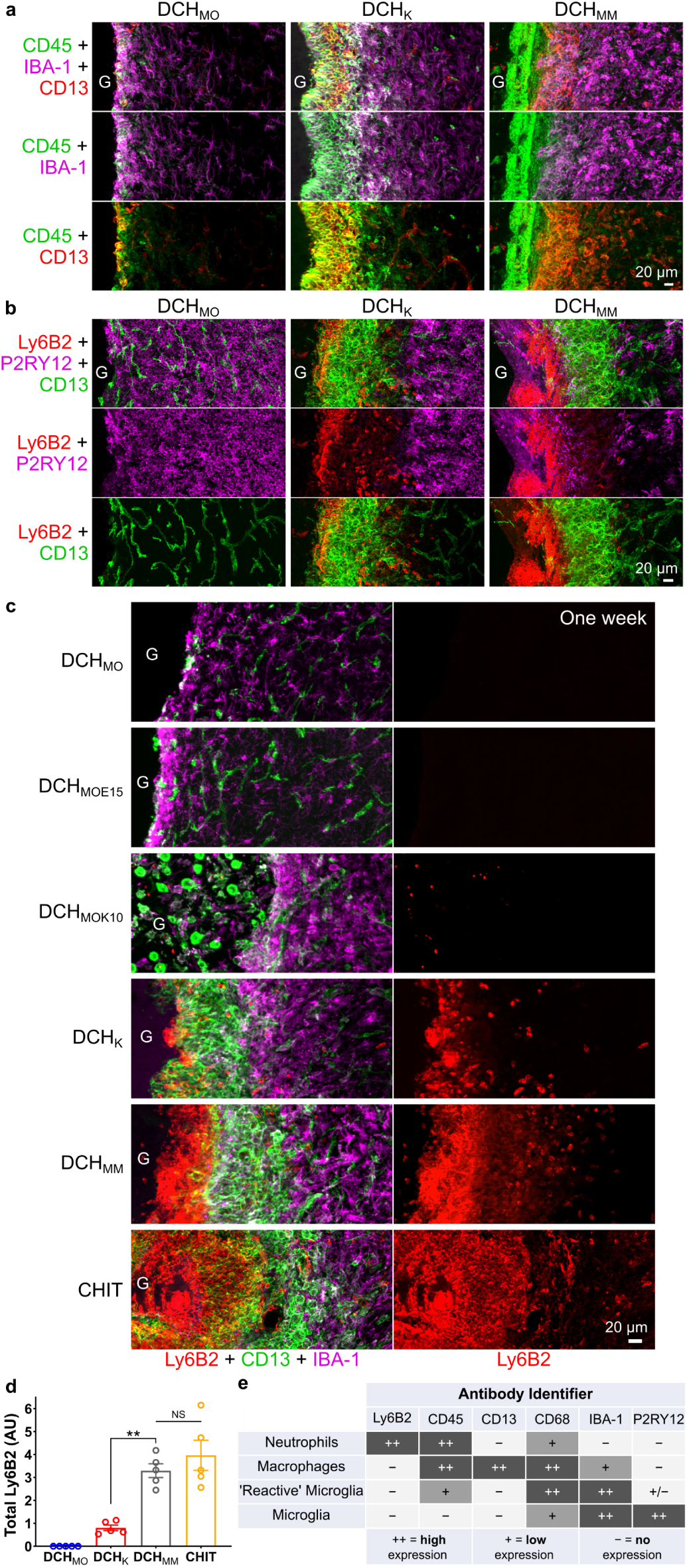
Different hydrogels attract inflammatory responses that differ in intensity and cellular phenotype. **a.** Detail images of differences in inflammatory cell recruitment one week after injection of DCH_MO_, DCH_K_, DCH_MM_ hydrogels (G). **b.** Detail images showing that CNS-derived P2RY12 positive microglia do not migrate into non-neural tissue compartments in spite of different FBRs evoked by DCH_MO_, DCH_K_, DCH_MM_. **c.** Detail images of hydrogel-tissue interface showing the escalating recruitment and persistence of Ly6B2-positive neutrophils at cationic hydrogel surfaces at one week after injection. **d.** Quantification of total Ly6B2 staining for hydrogels one week after injection. NS not significant and **P < 0.002 for hydrogels versus DCH_MM_, one-way ANOVA with Bonferroni. **e.** Table summarizing the combination of antibodies used to identify different inflammatory cell types. Graphs show mean ± s.e.m with individual data points showing *n* mice per group. (one column width figure)

**Fig. 4.**
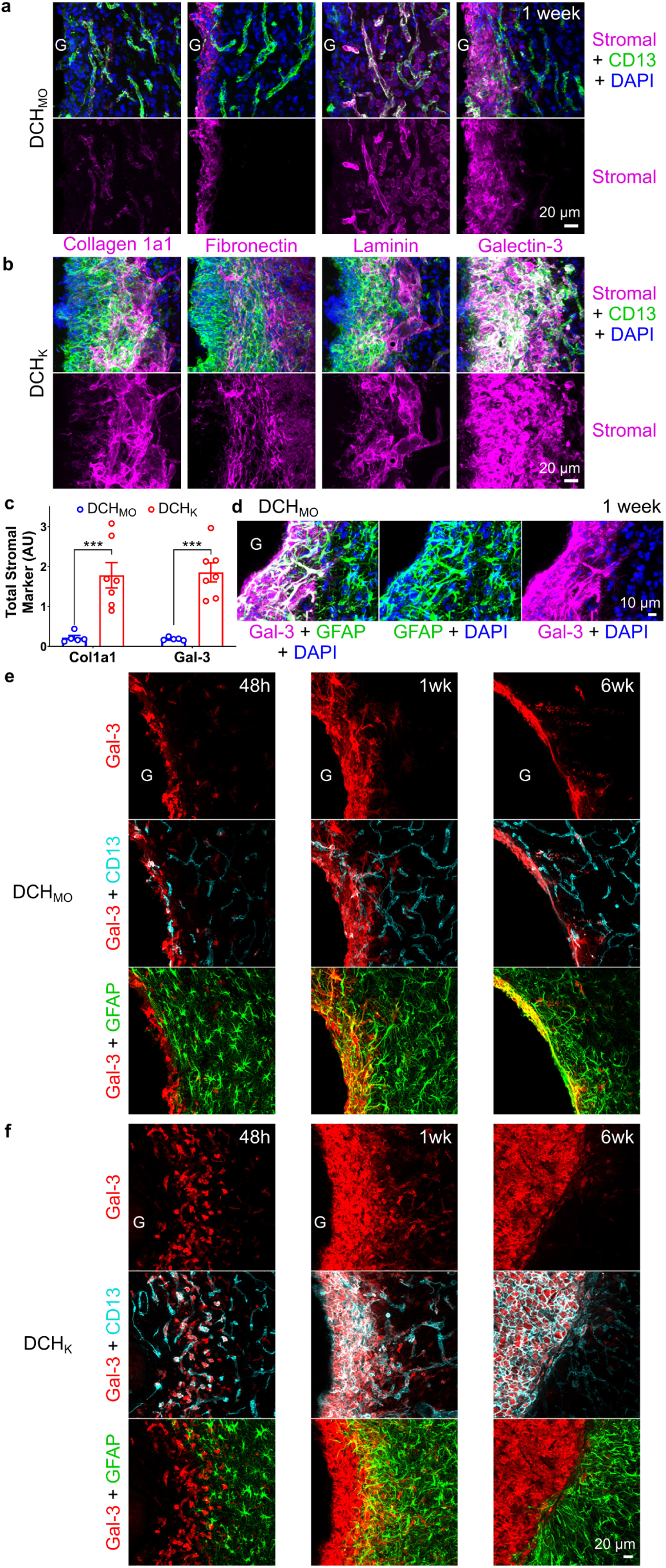
Hydrogels evoke fibrotic responses at the material-tissue interface that can involve stromal or astroglial cells. **a,b.** Detail images of stromal-associated markers, collagen 1a1 (Col1a1), fibronectin, laminin, and galectin-3 (Gal-3) and their relationship to non-neural CD13 positive cells at the tissue interface of DCH_MO_ (a) and DCH_K_ (b) at one week after injection (hydrogels, G). Colocalization of markers with CD13-positive cells is seen as white staining. **c.** Quantification of total Col1a1 and Gal-3 staining at one week after hydrogel injection. ***P < 0.0001 for DCH_MO_ versus DCH_K_ for the two stromal markers, two-way ANOVA with Bonferroni. Graphs show mean ± s.e.m with individual data points showing *n* mice per group. **d.** Detail image showing that adjacent to DCH_MO_, Gal-3 colocalizes with a narrow band of GFAP positive astrocytes that border the gel at 1 week. Gal-3 and GFAP colocalization is seen as white staining. **e,f.** Detail images showing the temporal evolution of Gal-3 expression at the interface of DCH_MO_ (e) and DCH_K_ (f) at acute (48 hours, 48h), subacute (1 week,1wk) and chronic (6 weeks, 6wk) timepoints after injection. Gal-3 and GFAP colocalization is seen as yellow staining. Gal-3 and CD13 colocalization is seen as white staining. (one column width figure)

**Fig. 5.**
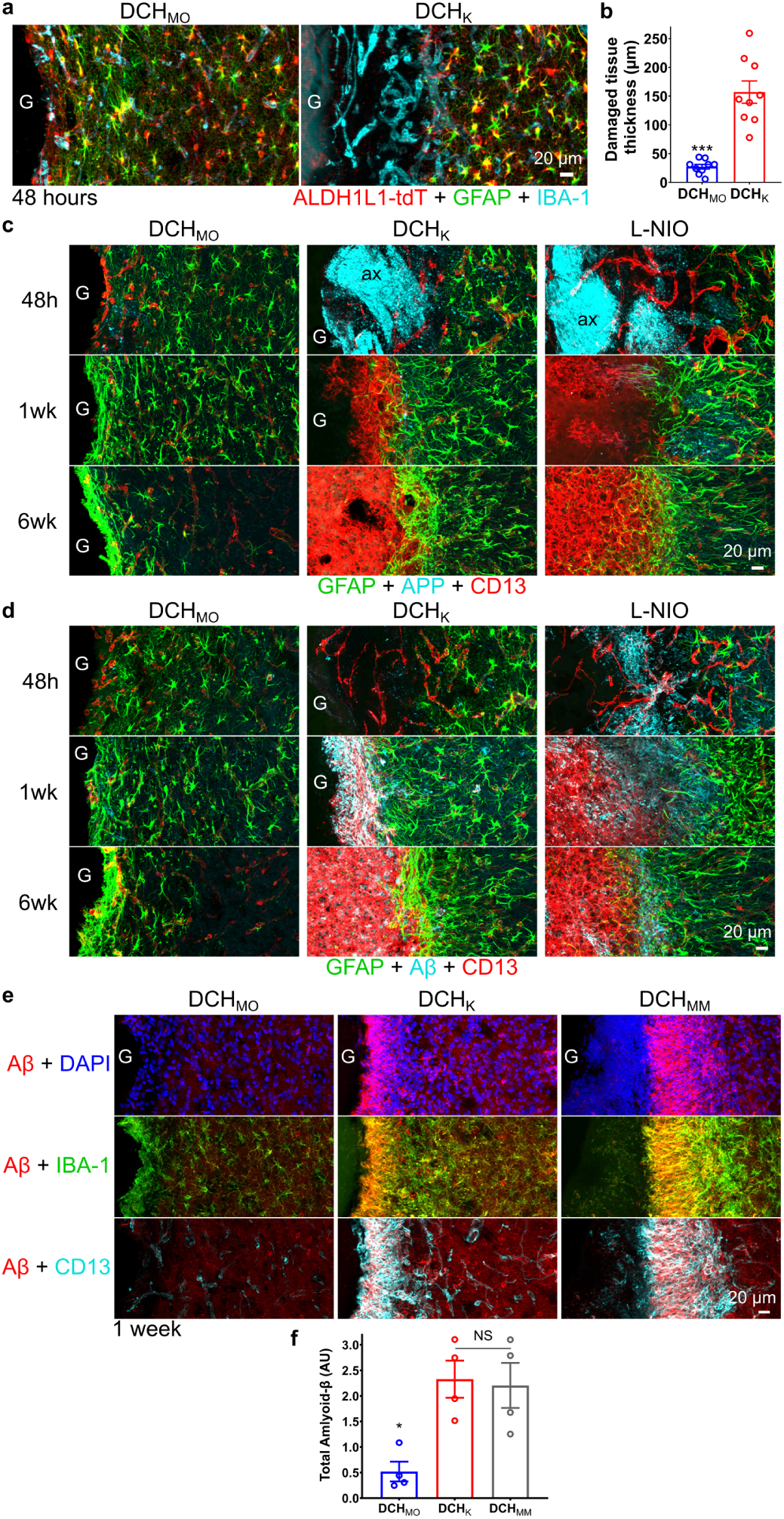
Hydrogel FBR severity is determined by the extent of acute neural tissue damage at the material interface, which is associated with axonal damage, APP accumulation and amyloid formation. **a.** Detail images of the material-tissue interface for DCH_MO_ and DCH_K_ at 48 hours after injection. ALDH1L1-tdT reporter and GFAP identify host astrocytes and the loss of these cells demarcates regions of host tissue damage. **b.** Quantification of radial thickness of tissue damage around DCH_MO_ and DCH_K_. ***P < 0.0001 Welch’s (unequal variance) t-test. **c,d.** Detail images for DCH_MO_, DCH_K_, and L-NIO-induced stroke at acute (48 hours, 48h), sub-acute (1 week, 1wk) and chronic (6 weeks, 6wk) timepoints after injection, comparing staining for amyloid precursor protein (APP) (c) or amyloid-beta (Aβ) (d) with GFAP and CD13. **e.** Images show Aβ, Iba-1 and CD13 at the interface of DCH_MO_, DCH_K_ and DCH_MM_ with host tissue at one week after injection. **f.** Quantification of Aβ at one week after DCH_MO_, DCH_K_ and DCH_MM_. NS or *P < 0.02 for hydrogels versus DCH_K_, one-way ANOVA with Bonferroni. Graphs shows mean ± s.e.m with individual data points showing *n* mice per group. G hydrogels, ax axons. (one column width figure)

Qualitative and quantitative examination showed that nonionic MC, like nonionic DCH_MO_, exhibited barely detectable levels of CD13-positive cells at interfaces with host tissue (Fig. 2a,c; Supplementary Fig. 4a). Moreover, anionic hydrogels HAMC as well as glutamate (E) inclusive DCH_E_ and DCH_MOE15_ also exhibited barely detectable levels of CD13-positive cells at host interfaces (Fig. 2a,c; Supplementary Fig. 4a). However, in striking contrast, cationic hydrogels, DCH_K_, DCH_MM_, DCH_MOK15_, DCH_MOK10_ and CHIT all exhibited large rims of CD13-positive cells infiltrating into deposits around their entire host interface and had significantly higher CD13-levels in central non-neural compartments, which did not differ in magnitude across different cationic hydrogels (Fig. 2b,c; Supplementary Fig. 4b). To directly probe the relationship between cationic charge and FBR, we incorporated variable amounts of lysine (K) into nonionic DCH_MO_ as a statistical copolymer [8] (Supplementary Fig. 3b). DCH_MOK15_ and DCH_MOK10_ both exhibited outer rims of infiltrating CD13 cells similar to DCH_K_, but with more cells infiltrating into the deposit center, suggesting that small amounts of dispersed cationic charge attracted CD13 cells, whereas dense charge in DCH_K_ prevented these same cells from distributing throughout the bulk of the material. In addition, we generated cationic, methyl-sulfonium-based DCH_MM_ (Supplementary Fig.3b) [8], which also exhibited a rim of infiltrating CD13 cells similar to DCH_K_, but in addition exhibited many DAPI-positive but CD13-negative cells that formed a distinct and separated layer of cells that directly interfaced with the hydrogel deposit surface (Fig. 2b). Similar CD13-negative cells at the material interface were not obviously present in other DCH, but were present within CHIT deposits (Fig. 2b), suggesting a distinct cellular infiltrate requiring further characterization as conducted below.

These findings show that hydrogel FBRs vary with definable hydrogel properties and implicate cationic charge presented at material interfaces with host tissue as a major factor determining FBR severity. Remarkably, despite significantly elevated CD13 in the non-neural compartments of cationic hydrogels (Fig. 2a-c), there was no significant difference in either CD13 (Fig. 2d; Supplementary Fig. 5) or GFAP (Fig. 2e) levels, and no detectable qualitative difference in NeuN-positive neuron density (Supplementary Fig. 5), in the neural tissue adjacent to either cationic, nonionic or anionic hydrogels, suggesting that neural tissue beyond the interface between hydrogel and host was not substantively differentially impacted. The unexpected finding that GFAP levels were indistinguishable in neural tissue adjacent to hydrogels that evoked markedly different FBR at host interfaces indicates that GFAP staining is not a sensitive or sufficient marker with which to evaluate and differentiate FBRs.

### Hydrogel FBRs differ in inflammatory cell recruitment/persistence

To discriminate among diverse innate-immune inflammatory cells involved in hydrogel FBRs, we used immunohistochemistry for: (i) CD45 to identify leukocytes and reactive CNS microglia [35]; (ii) Iba-1 to identify blood-borne monocytes/macrophages and CNS microglia [35]; (iii) P2RY12 to identify only microglia [20, 36, 37]; (iv) Ly6B2 to identify neutrophils [21]; and (v) CD13 to identify myeloid lineage peripheral inflammatory cells such as monocytes and macrophages or fibroblast lineage cells (Fig. 3) [28]. We examined combinations of these markers at one week after hydrogel injections (Fig. 1b).

For all hydrogels, the neural tissue immediately adjacent to deposits contained reactive, CNS-derived microglia that were Iba-1-, CD45- and P2RY12-positive and CD13-negative that intermingled with astrocytes and neurons, remained within the neural tissue compartment and did not detectably infiltrate into any hydrogel (Figs. 3a,b; Supplementary Fig.6a). P2RY12 expression was reduced in reactive microglia that dispersed directly within forming astroglia borders. Immediately abutting these microglia, hydrogel deposits exhibited variable intensities of inflammatory cell infiltrates that were CD45- and CD13-positive, and P2RY12-negative, and were peripherally-derived myeloid lineage leukocytes and not CNS microglia (Fig. 3a,b, Supplementary Fig. 6a,b). A comparable segregation of blood-borne inflammatory cells into a non-neural tissue compartment away from neural tissue was also observed for the L-NIO stroke lesion (Supplementary Fig. 6c).

**Fig. 6.**
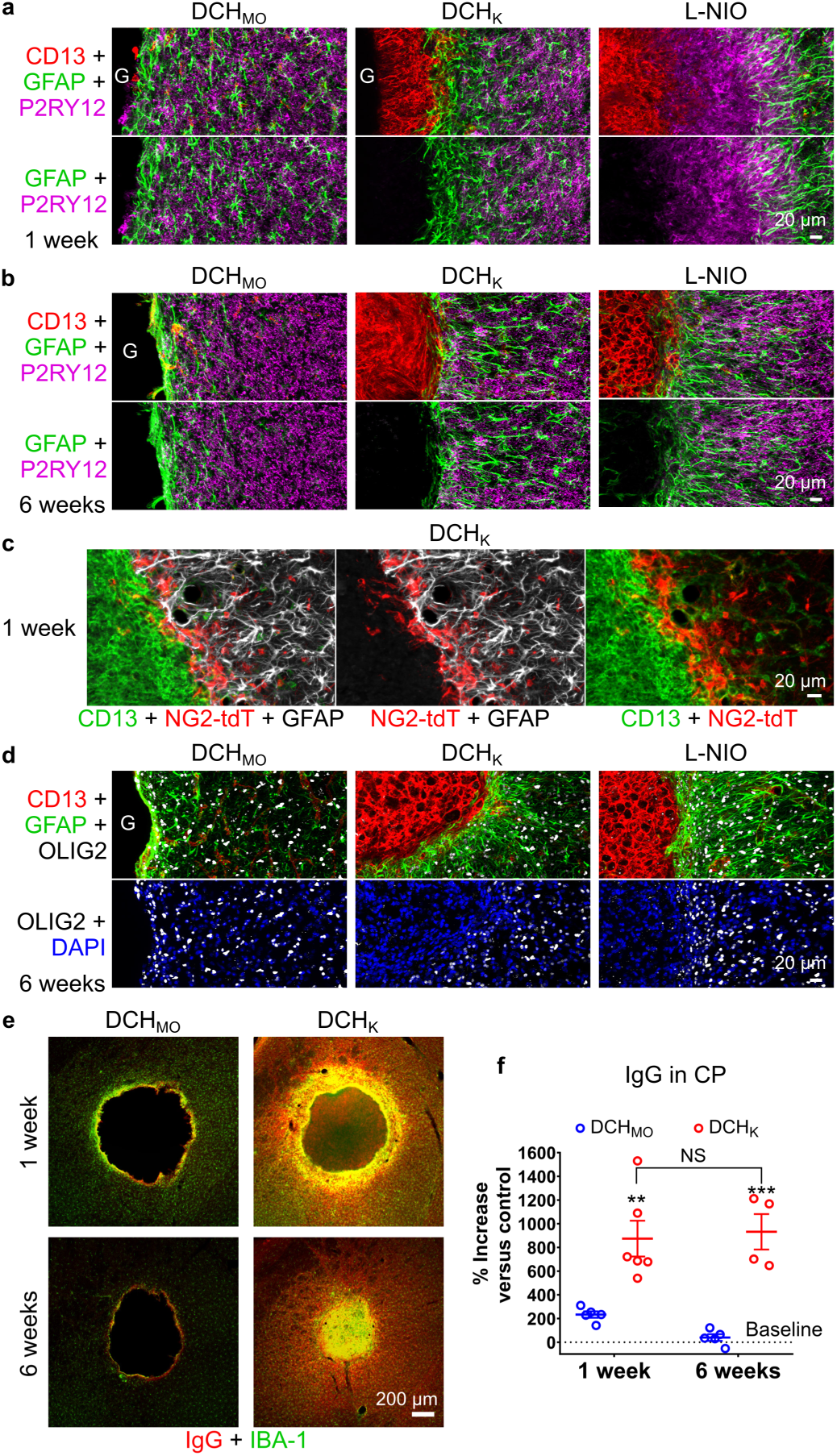
Astrocytes form limitans borders that isolate hydrogels and non-neural FBR components from viable neural tissue. **a,b.** Progression of astrocyte limitans border formation from 1 week (a) to 6 weeks (b) for DCH_MO_, DCH_K_, and L-NIO-induced stroke. **c,d.** OPC identified by NG2-targeted reporter (tdT) (c) and Olig2 (d) intermingle with GFAP-positive astrocytes and do not migrate into CD13-positive regions. **e.** Comparison of extent of blood brain barrier (BBB) disruption and repair as measured by IgG staining around DCH_MO_ and DCH_K_ deposits at 1 and 6 weeks after injection. **f.** Quantification of the percentage increase in IgG levels in the hydrogel injected caudate putamen (CP) normalized to the non-injected contralateral side. **P < 0.001 and ***P < 0.0001 for DCH_K_ versus DCH_MO_ at 1 week and 6 week respectively and NS for DCH_K_ samples between the two time points, two-way ANOVA with Bonferroni. Graph shows mean ± s.e.m with individual data points showing *n* mice per group. (one column width figure)

Different hydrogels exhibited remarkable variation in both the intensity and cellular phenotype of the blood-borne inflammatory response that they attracted. Nonionic DCH_MO_ attracted only rare infiltrating CD45-positive leukocytes at interfaces with host tissue, and these were macrophage lineage (Iba1-positive and CD13-positive) (Fig. 2a,3a,b; Supplementary Fig. 6a). In contrast, cationic DCH_K_ and DCH_MM_ both attracted thick rims of infiltrating macrophage-lineage cells (CD45-positive, Iba1-positive and CD13-positive) along their entire host interfaces (Fig. 2b,3a,b; Supplementary Figs. 4,6a). Immunohistochemical staining also revealed that the thick layers of DAPI-positive and CD13-negative cells infiltrating into DCH_MM_ and CHIT (Fig. 2b) were leukocytes (CD45-positive), but were not macrophage lineage (Iba1-negative) (Fig. 3a), and instead were highly phagocytic and cytotoxic Ly6B2-positive neutrophils (Fig. 3b,c; Supplementary Fig. 6b). In striking contrast, nonionic DCH_MO_ and weakly cationic DCH_MOK10_ or anionic DCH_MOE15_, contained no detectable Ly6B2-positive neutrophils, while strongly cationic DCH_K_ attracted a few of these cells (Fig. 3c,d; Supplementary Fig. 6a).

To test whether inflammatory response intensity correlated directly with strength of cationic charge at hydrogel surfaces, we measured zeta potentials of DCH hydrophilic polymer chains. DCH_K_, containing homopolymers of lysine, had a significantly greater positive charge at physiological pH than DCH_MM_ (containing statistical copolymers with 10% uncharged alanine) due to a higher total number of charged side chain groups along each hydrophilic chain (Supplementary Fig. 6d). However, DCH_K_ had significantly lower levels of neutrophils and equivalent levels of macrophages compared to DCH_MM_ (Fig. 3d). Nonionic DCH_MO_ had no detectable charge, no infiltration of neutrophils and only rare macrophages (Fig. 3a,b,c; Supplementary Fig. 6d).

Persistence of peripheral inflammatory cells at hydrogel deposits was evaluated at three weeks after injection (Supplementary Fig. 7a,b). Notably, while Ly6B2-positive neutrophils were mostly resolved at CHIT deposits by 3 weeks, thick rims of these acute inflammatory cells were sustained at DCH_MM_ deposits with a distribution that was not obviously different to that at one week. DCH_MM_ deposits also remained largely un-resorbed at 3 weeks whereas CHIT injections were completely infiltrated by cells at this time point (Supplementary Fig. 7c).

**Fig. 7.**
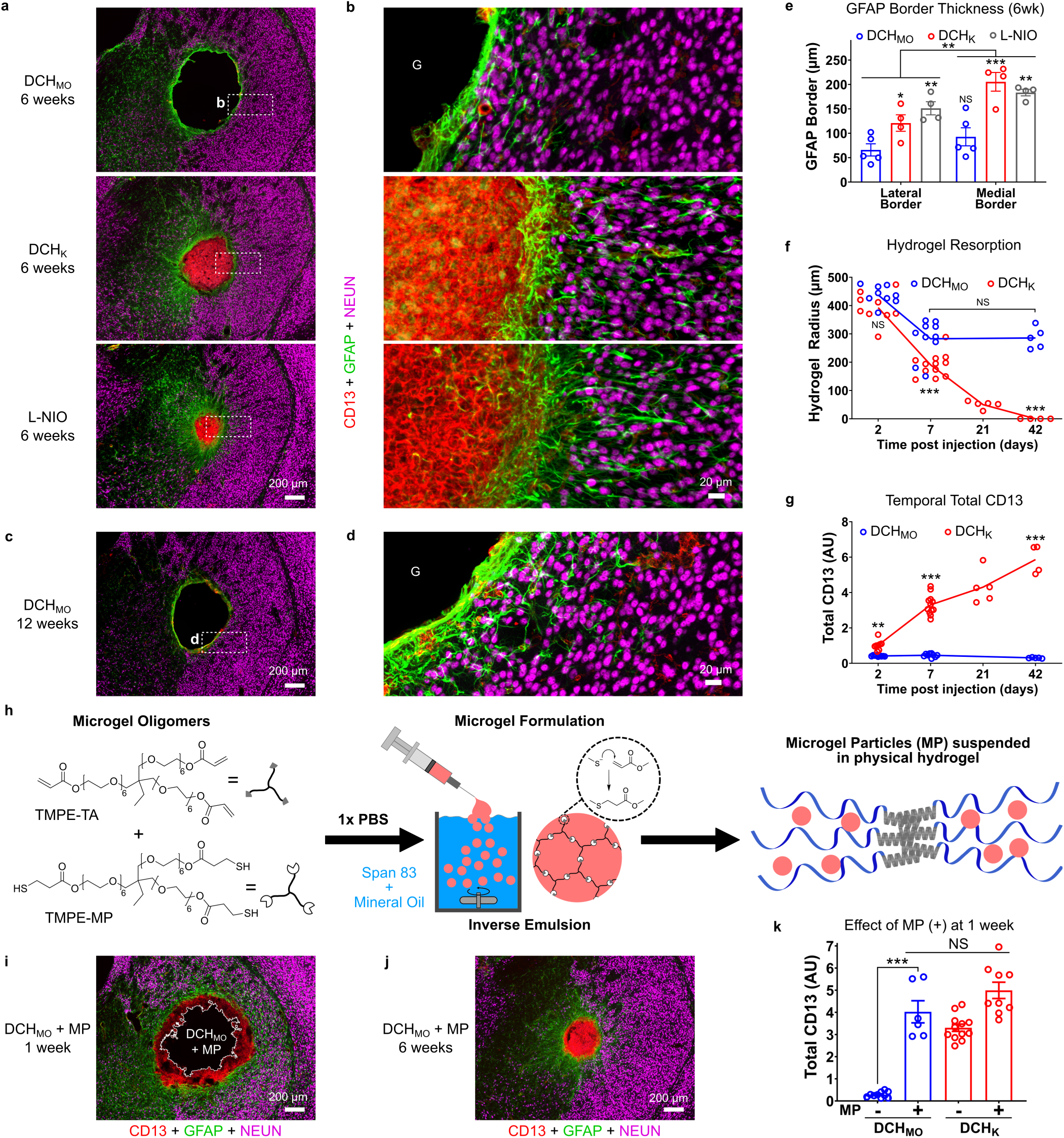
Hydrogel FBR with recruitment of CD13 positive cells leads to hydrogel resorption. **a,b.** Survey and detail images show qualitative reductions in size of DCH_K_ deposit and L-NIO-induced infarct, but not of DCH_MO_ deposit at 6 weeks after injection **c,d.** Survey and detail images show DCH_MO_ deposit remains largely unresorbed up to 12 weeks after injection. **e.** Quantification of GFAP positive cell border thickness (measurement of half width maximum of GFAP intensity curve) at the lateral (grey matter) or medial (internal capsule) borders at 6 weeks. *P<0.05 and **P < 0.003 and ***P < 0.0001 versus DCH_MO_ on same side; NS between DCH_MO_ on either side, **P < 0.003 for border location effect across all samples, two-way ANOVA with Tukey. **f.** Quantification of change in hydrogel radius for DCH_MO_ and DCH_K_ from 2 to 42 days. NS, **P < 0.003 and ***P < 0.0001 for DCH_K_ versus DCH_MO_ at 2, 7 and 42 days; NS for DCH_MO_ at 7 versus 42 days, two-way ANOVA with Bonferroni. **g.** Quantification of change in total CD13 for DCH_MO_ and DCH_K_ from 2 to 42 days. **P < 0.007 and ***P < 0.0001 for DCH_K_ versus DCH_MO_ at the same timepoints. NS between the various time points for DCH_MO_, two-way ANOVA with Bonferroni. **h.** Schematic of microgel particle (MP) synthesis involving the inverse emulsion of polyethylene glycol (PEG)-based thiol and acrylate functionalized oligomers that react via Michael addition. MP can be readily suspended in physical hydrogels tested. **i,j.** Survey images of FBR to DCH_MO_ loaded with MP at 1 week and 6 weeks show that incorporation of MP into DCH_MO_ leads to recruitment of CD13 positive cells and hydrogel resorption. **k.** Quantification of the effect of incorporation of MP (+) on total CD13 positive cell response for DCH_MO_ and DCH_K_ at 1 week. NS and ***P < 0.0001, two-way ANOVA with Bonferroni. All graphs show mean ± s.e.m with individual data points showing *n* mice per group. (two column width figure)

These findings show that the intensity and duration of the inflammatory cell components of CNS FBRs varied markedly in response to deposits of different hydrogels and that cationic charge at material-host interfaces is an important factor in attracting blood-borne phagocytic leukocytes. Nevertheless, charge magnitude alone does not determine the nature or distribution of phagocytes attracted.

### Hydrogel FBRs involve stromal cells and fibrosis

To characterize fibrosis associated with hydrogel FBRs, we compared a representative hydrogel with limited detectable CD13 levels (DCH_MO_) to a representative hydrogel with a pronounced but focal rim of CD13 cells (DCH_K_) using immunohistochemistry for cell surface and extracellular matrix molecules, collagen-1a1, fibronectin, laminin, and galectin-3 with staining for CD13 at various times after hydrogel injections (Figs. 1b,4a-e). Collagen-1a1, fibronectin and laminin demarcate stromal cells [38], whereas galectin-3 can demarcate multiple cell types including stromal and inflammatory cells [39–41]. In uninjected normal neural tissue, (i) CD13 stained stromal cells along blood vessels and at the meninges, (ii) collagen-1a1 and laminin lightly decorated cell surfaces along blood vessels and around neurons, (iii) fibronectin was present at large vessels and at the meninges, and (iv) galectin-3 was essentially undetectable [42] (Supplementary Fig. 8a).

**Fig. 8.**
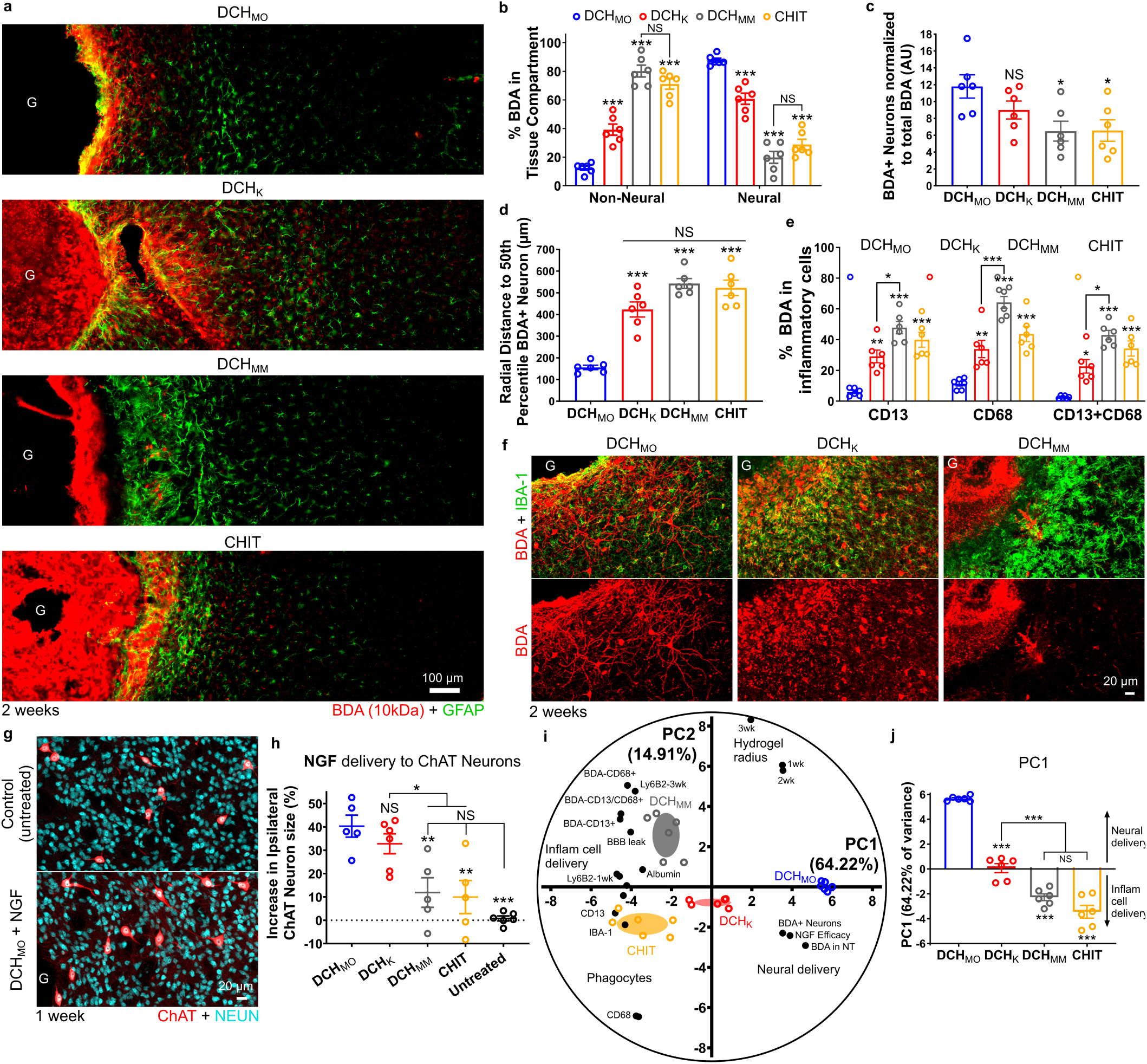
Hydrogel FBR alters CNS Molecular delivery. **a.** Detail images of BDA (10kDa) biodistribution released from DCH_MO_, DCH_K_, DCH_MM_, and CHIT at 2 weeks. **b.** Quantification of BDA in tissue compartments adjacent to hydrogels. ***P < 0.0001 versus DCH_MO_; NS not significant **c.** Quantification of the normalized number of BDA-positive neurons at 2 weeks. *P < 0.05, NS versus DCH_MO_. **d.** Quantification of the radial distance to the 50% percentile BDA-positive neuron from the hydrogel-tissue interface. ***P < 0.0001 versus DCH_MO_. **e.** Quantification of BDA-positive inflammatory cells at 2 weeks. *P < 0.005, **P < 0.001, ***P < 0.0001 versus DCH_MO_. **f.** Detail images showing phenotype of BDA positive cells adjacent to hydrogel deposits. **g.** Detail images showing increase in cholinergic (ChAT-positive) neuron size in striatum by NGF delivered from DCH_MO_. **h.** Quantification of the increase in cholinergic neuron size normalized to the contralateral side at 1 week after injection of NGF releasing hydrogels. ***P < 0.0001 and **P < 0.005 versus DCH_MO_, NS for DCH_MM_ and CHIT versus untreated, *P < 0.05 for DCH_MM_ and CHIT versus DCH_K_, NS between DCH_MO_ and DCH_K_. **i.** Principal Component Analysis (PCA) for DCH_MO_, DCH_K_, DCH_MM_, and CHIT hydrogels incorporating data from molecular delivery and immunohistochemistry evaluations. **j.** Graphical representation of the positions of hydrogels along PC1 axis (accounting for 64.22% of total variance) with the positive direction representing effective neural molecular delivery while the negative direction represents molecular delivery to inflammatory cells, ***P < 0.0001 for all versus DCH_MO_, ***P < 0.0001 for DCH_K_ versus DCH_MM_ and CHIT, NS for DCH_MM_ versus CHIT. For all graphs, data are mean ± s.e.m with individual data points superimposed over bar graphs showing *n* mice per group. Statistical analysis using one or two-way ANOVA with Bonferroni where appropriate. (two column width figure)

At one week after injection, tissue adjacent at interfaces with non-ionic DCH_MO_ exhibited staining for collagen-1a1 and laminin only along blood vessels in a manner similar to normal neural tissue, whereas fibronectin and galectin-3 were moderately elevated in narrow rims along host-gel interfaces (Fig. 4a,c-e). In contrast, the thick capsule of CD13-positive cells circumscribing cationic DCH_K_ exhibited significantly greater levels of collagen-1a1, laminin, fibronectin and galectin-3 (Figs. 4b,c). Around DCH_K_, galectin-3 was robustly expressed by essentially all CD13-positive cells, whereas collagen-1a1, laminin and fibronectin demarcated subsets of stromal cells in outer margins (Fig. 4b) that interfaced directly with the forming astrocyte limitans border. An interior layer of CD13- and galectin-3-positive but collagen-1a1, fibronectin and laminin-negative cells, likely peripheral inflammatory cells, interacted directly with the DCH_K_ surface (Fig. 4b). This inflammatory and fibrotic cell stratification around DCH_K_ is reminiscent of the cellular organization in CNS abscesses and traumatic injury lesions [40, 43]. CD13- and galectin-3-positive cells persisted chronically in the non-neural tissue compartment of resorbed DCH_K_ for at least 6 weeks (Fig. 4f). Interestingly, the GFAP-positive astrocyte limitans border that formed the direct host interface with non-ionic DCH_MO_ also extensively expressed galectin-3 and this expression persisted chronically for at least 6 weeks (Fig. 4d,e; Supplementary Fig. 8b).

These findings show that DCH_MO_ evokes minimal levels of fibrosis, whereas cationic DCH_K_ evokes a stronger fibrotic FBR with elevated levels of multiple matrix molecules at the interface with preserved neural tissue. Additionally, we identified a possible conserved function for galectin-3 in isolating foreign bodies from parenchymal neural tissue regardless of the severity of the FBR, with its expression occurring rapidly following hydrogel injection and present only in cells forming the material-tissue interface, regardless of cell type (inflammatory cells, stromal cells or astrocytes). Galectin-3 is known to be associated with fibrosis and wound healing in various tissue injuries including kidney, liver and heart [44] and is significantly upregulated in macrophages/microglia and reactive astrocytes after various traumatic CNS injuries [42, 45–47]. Here we identify galectin-3 to also be an important constituent of CNS FBRs to materials.

### Tissue damage drives FBR and causes acute amyloid formation

We next looked for associations between FBR, inflammation, fibrosis and host tissue loss, and for other potential molecular features shared across FBR and CNS wound response. To do so, we first used mice expressing reporter protein transgenically-targeted to host astrocytes via Aldh1l1-Cre-ERT2 and evaluated host neural tissue damage and changes in various molecular markers, including accumulation of amyloid precursor protein (APP), a robust marker of axonal injury [48] and one of its breakdown products, beta-amyloid (Aβ) [49]. We compared representative hydrogels displaying minimal or pronounced FBRs with the CNS wound response to L-NIO stroke lesions at various time points.

At 48 hours after injection, DCH_MO_ exhibited a minimal zone of host neural tissue loss averaging only 25µm, the equivalent of one or two cells in thickness, whereas DCH_K_ exhibited a significantly greater, but still relatively small zone of host neural tissue loss of about 150µm in thickness, as evidenced by: (i) astrocyte and neuron loss but persistence of vasculature, (ii) presence and phagocytosis of reporter protein debris, (iii) extravasation of blood borne fibronectin [50], (iv) up-regulation of CD13 expression along thin PECAM-1-positive blood vessels within the damaged tissue zone and (v) infiltration of blood-borne phagocytic leukocytes (Fig. 5a,b; Supplementary Fig. 9a-e).

**Fig. 9.**
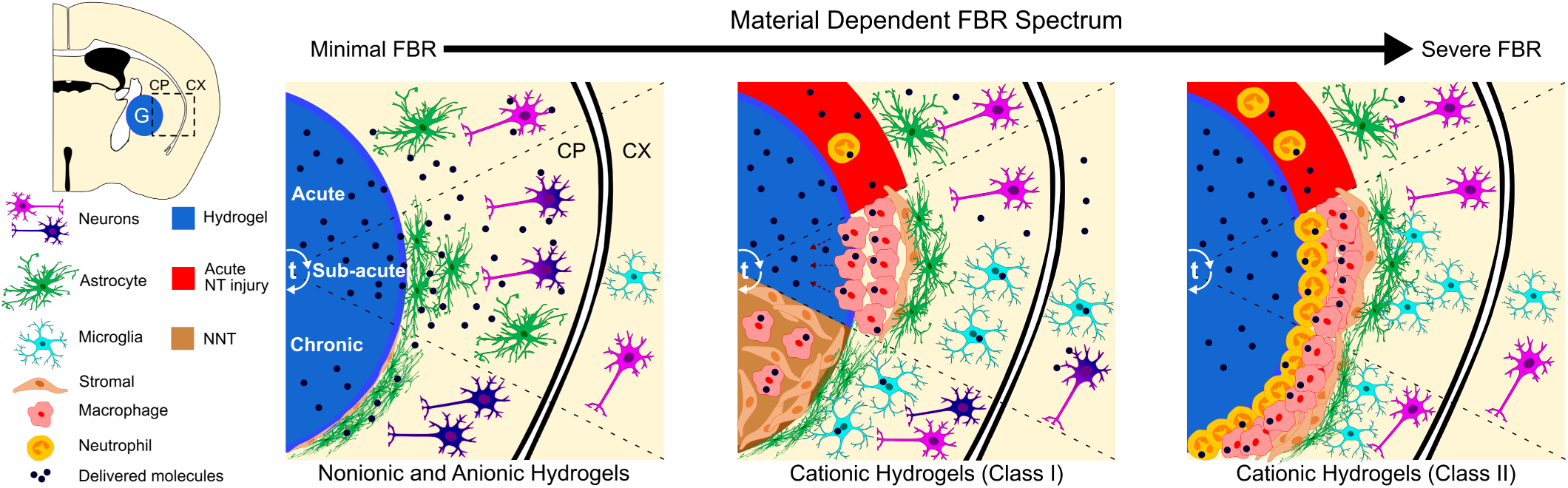
Schematic summarizing the main findings of this study: CNS FBRs mimic the natural wound healing response to CNS tissue injury as it progresses over time (t) from acute to chronic stages (top to bottom). CNS FBRs to biomaterials exist on a severity spectrum (left to right) that is dependent on definable material properties such as the biomaterial chemistry presented to the host at the biomaterial-tissue interface. CNS FBR phenotype and its severity alter hydrogel function in terms of molecular delivery and biomaterial resorption. (two column width figure)

Further characterization of the extent of axonal injury was provided by amyloid precursor protein (APP) staining. APP is low or undetectable by immunohistochemistry in healthy CNS tissue, but increases markedly after axonal injury [48, 51]. After 48 hours, DCH_MO_ displayed only small amounts of APP accumulation at the hydrogel-host interface (Fig. 5c). In contrast, APP was present throughout the narrow zone of neural tissue damage induced by DCH_K_ and APP accumulation was prominent in damaged axon bundles close to DCH_K_ deposits in a manner comparable with L-NIO stroke lesions (Fig. 5c). By one week, APP staining had decreased substantially and was detectable only at the interface of neural to non-neural tissue, and APP had returned to baseline at six weeks across all conditions (Fig. 5c).

The formation of beta-amyloid (Aβ) via cleavage of APP by β- and γ-secretase enzymes is increasingly recognized as a normal occurrence during CNS wound responses [52, 53]. We evaluated the progression of APP to Aβ (Fig. 5d). At 48 hours, Aβ was sparse at the hydrogel-tissue interface for both DCH_MO_ and DCH_K_, whereas L-NIO stroke lesions already showed clear Aβ accumulation at the borders of infarcted and spared neural tissue (Fig.5d). By one week, Aβ was barely detectable around DCH_MO_ and was comparable to PBS injections where Aβ was constrained to the track of tissue disrupted by the injection pipette (Supplementary Fig. 10a,b). At one week after DCH_K_, DCH_MM_ injections and L-NIO stroke lesions, Aβ accumulation was most prominent within the layers of CD13-positive inflammatory cells that infiltrated areas of neural tissue damage (Fig. 5d,e). In addition, in all conditions, there was prominent colocalization of Aβ within Iba-1-positive reactive microglia in adjacent spared neural tissue, suggesting phagocytosis by these cells as well (Fig. 5e, Supplementary Fig. 10c). Quantification confirmed significantly higher levels of Aβ at one week for DCH_K_ and DCH_MM_ compared to DCH_MO_ (Fig. 5e,f). Increased neutrophil accumulation and persistence in DCH_MM_ compared to DCH_K_ (Fig. 3c,d) was not associated with any increase in Aβ formation (Fig. 5e,f) suggesting that Aβ is formed as a result of acute neural tissue injury and not by ongoing chronic inflammation. At 6 weeks, Aβ levels had reduced markedly and were evident only within the persistent non-neural tissue lesions for DCH_K_ and L-NIO stroke (Fig. 5d).

These finds show that the inflammatory and fibrotic FBRs evoked by cationic hydrogels were driven by narrow but measurable zones of host neural tissue damage occurring along hydrogel-host interfaces soon after injection and that this was essentially absent with the nonionic DCH_MO_. These findings also show that APP accumulation and Aβ formation around hydrogel deposits occurred as a result of damage to neural tissue and axonal injury. These observations are consistent with growing evidence that APP accumulation and subsequent Aβ formation are part of a conserved innate wound and foreign body response. While the specific functions of Aβ production in this context are not yet defined, recent evidence suggests that Aβ may exert antimicrobial activities [52, 53].

### Astrocyte limitans borders isolate FBRs from neural tissue

Neural tissue contains several types of glial cells that become reactive around damaged CNS tissue and participate in FBRs. Astrocytes are well known to form ‘scars’ that serve as limitans borders to isolate CNS lesions [19]. Microglia contribute to border formation and exert phagocytic functions [20] while oligodendrocyte progenitors (OPC) proliferate and repair myelin [22]. We identified astrocytes and microglia by immunohistochemistry for GFAP and P2RY12 respectively, and OPC by staining for *Olig2* and the reporter protein, td-Tomato (tdT), which had been transgenically-targeted by NG2(*Cspg4*)-Cre-ERT2 [54]. We compared combinations of markers and probed blood-brain barrier (BBB) integrity at one and six weeks after hydrogel injection or L-NIO stroke (Figs. 6,7).

GFAP-positive astrocytes had begun to form distinct borders around all hydrogel deposits by one week after injections, and these borders persisted and consolidated by six weeks. Astrocyte borders around hydrogels were similar in appearance to borders formed around L-NIO infarcts and clearly segregated persisting hydrogel material and CD13-positive stromal and inflammatory cells from neural tissue containing NeuN-positive neurons (Figs. 1c,d;2a,b;6a,b;7a,b). Notably, astrocyte borders adjacent to DCH_MO_ and other nonionic hydrogels interfaced directly with gel surfaces with limited intervening fibrosis or inflammation, whereas astrocyte borders adjacent to DCH_K_ and other cationic gels or L-NIO interfaced with stromal and inflammatory cells (Figs. 2a,b,4,6a,b). Astrocyte borders around hydrogels and L-NIO infarcts were similar in appearance to astrocyte limitans borders that separate healthy neural tissue from non-neural stromal cells along all interfaces of normal CNS with meninges (Supplementary Fig. 1c) [55]. Astrocyte border thickness at six weeks was proportionally greater with increasing severity of FBR, such that DCH_MO_ displayed the thinnest astrocyte borders while DCH_K_ and L-NIO both had similar sized borders of more than double the thickness (Fig. 7e). Across all conditions at six weeks, lateral astrocyte borders formed by grey matter astrocytes, were thinner compared to medial borders that recruited white matter astrocytes from the internal capsule (Figs. 1b,7a-b,e).

Neural parenchyma extending from astrocyte borders exhibited initially moderate astrocyte reactivity indicated by elevated GFAP, which declined significantly over time and by six weeks was minimal in all cases (Fig. 6a,b;7a,b). Reactive microglia and OPC intermingled with astrocytes along borders and adjacent neural tissue, but did not infiltrate into hydrogels or contribute to volumes of CD13-positive fibrosis and inflammation at any timepoint examined (Fig. 6a-d and Supplementary Fig. 11a). At six weeks a thin, single cell layer of non-neural tissue (CD13-positive) that expressed stromal cell markers PDGFRβ and fibronectin interfaced directly with DCH_MO_ and the astrocyte limitans border (Supplementary Fig. 11b) while these cells contributed to large volumes of non-neural fibrotic tissue in DCH_K_ and L-NIO strokes (Supplementary Fig. 11b,c).

Astrocyte border formation is essential for re-establishing BBB integrity after CNS injuries [30]. To probe BBB integrity adjacent to hydrogel deposits, we stained for mouse immunoglobulin (IgG) and albumin, the two most abundant serum proteins [30]. As expected, after one week and prior to border formation [30], IgG and albumin staining were increased in neural parenchyma around hydrogel injections and L-NIO stroke lesions, and were significantly higher around DCH_K_, compared with DCH_MO_ (Fig. 6e,f; Supplementary Fig. 12a-c). By six weeks, serum protein levels around DCH_MO_ were indistinguishable from those in uninjured tissue and were restricted to a thin layer of non-neural tissue that interfaced with the astrocyte limitans border. In contrast, around L-NIO stroke lesions and cationic DCH_K_, serum proteins remained significantly elevated in neural tissue at six weeks (Fig. 6e,f, Supplementary Fig. 12a-c).

These findings show that astrocytes rapidly form limitans borders around hydrogels in a manner similar to borders formed around ischemic or traumatic tissue damage, or that exist along meningeal non-neural stromal tissue around healthy CNS. Astrocytes, microglia and OPCs become reactive in neural tissue adjacent to hydrogel deposits or stroke lesions, but do not migrate into the non-neural fibrotic tissue compartments that persist chronically. Glial reactivity and BBB leakiness into neural tissue adjacent to astrocyte borders persist longer adjacent to cationic materials that generate substantial inflammation and fibrosis compared with nonionic materials that do not.

### FBR determines hydrogel resorption or persistence

To evaluate the relationship between FBR and hydrogel resorption or persistence, we compared nonionic DCH_MO_ and cationic DCH_K_ deposits at various times after injection (Figs. 1,7). Quantification showed that after 48 hours, DCH_MO_ and DCH_K_ exhibited deposits of similar size. After one week, DCH_MO_ deposits had decreased in size by 30%, but remained at this size after six weeks and persisted essentially unchanged after twelve weeks (Figs. 7a-d,f). In contrast, after one week DCH_K_ deposits had decreased in size significantly by over 50% and continued to steadily decrease in size until there was no detectable deposit remaining at 6 weeks (Fig. 7a,b,f). Over time, DCH_MO_ exhibited no increase in CD13 levels above baseline, whereas DCH_K_ evoked a steadily increasing infiltration of CD13-positive inflammatory and stromal cells that proceeded in a concentric fashion from hydrogel-tissue interfaces inwards until the entire deposit was consumed (Figs. 1c,d,3a,4b,7a,b,g; Supplementary Fig. 7a). By comparison, the wound response to L-NIO-induced ischemia attracted a pronounced infiltration of CD13-positive cells that rapidly filled the entire volume of damaged tissue and then persisted (Figs. 1c,d,7a,b). These findings show that DCH_MO_ and other nonionic hydrogels persist and are not efficiently resorbed *in vivo* because they do not attract sufficient CD13-positive phagocytes. In contrast, DCH_K_ and other cationic hydrogels attract these phagocytic leukocytes to their interfaces with host tissue and are gradually resorbed from the outside in and become replaced by fibrosis. DCH_MM_ was an exception to this general cationic hydrogel resorption trend and instead showed minimal hydrogel resorption after 3 weeks (Supplementary Fig. 7c). This chronic hydrogel persistence coupled with the unresolved Lys6B2-positive neutrophils at the material-tissue interface suggests that DCH_MM_ is unable to be readily cleared due to frustrated phagocytosis. Thus, hydrogel properties determine FBR features, which in turn determine hydrogel resorption or persistence.

### Microparticles alter FBR and hydrogel resorption

Incorporating nano/microparticles into hydrogels may be a useful tool to enhance the control of delivery of multiple and diverse molecular cargos independently from a single construct to the CNS [18, 56, 57]. Since introduction of particles may alter the physiochemical properties of hydrogels into which they are loaded, we examined the effects on FBR and resorption of hydrogels laden with PEG based nonionic microparticles (MP) formulated via a standard inverse emulsion thiol-ene Michael addition process (Fig. 7h) [58–60]. MP, with an average diameter of 3.3 µm, imparted a modest increase in mechanical properties to DCH but did not alter injectability (Supplementary Fig. 13a-f). MP at high concentrations were non-toxic to neural progenitor cells *in vitro* (Supplementary Fig. 13d). As described above, nonionic DCH_MO_ and HAMC on their own evoked minimal fibrotic or inflammatory responses and formed long persisting deposits. Addition of nonionic MP to these hydrogels caused pronounced recruitment of peripheral inflammatory cell infiltration in the form of both CD13-positive macrophages and Lys6B2-positive neutrophils, as well as rapid hydrogel resorption and fibrotic replacement (Figs. 7i-k; Supplementary Fig. 13g-k). Nevertheless, the FBR of MP loaded DCH_MO_ did not detectably increase acute neural tissue loss or axonal injury (Supplementary Fig. 13g,h). Similar sized non-toxic MP as the ones evaluated here (~3 µm diameter) have previously demonstrated a high susceptibility to phagocytosis *in vitro* by macrophages [61]. Incorporating MP into nonionic hydrogels may stimulate similar size recognition programs *in vivo* thus leading to the recruitment of phagocytes by mechanisms other than those associated with host neural tissue damage at the hydrogel-host interface as was the case for cationic hydrogels. These findings show that adding such MP to hydrogels that would otherwise be ignored by phagocytes can induce a phagocytic FBR that steadily resorbs the material and replaces it with fibrosis.

### FBR alters hydrogel molecular delivery to CNS parenchyma

To characterize FBR effects on delivery of molecular cargos, we first examined the biodistribution of model non-bioactive molecules released into neural tissue adjacent to hydrogel deposits, and second evaluated the efficacy of bioactive growth factor delivery. As non-bioactive molecules, we used biotinylated dextran amines (BDA) of different molecular weights because they are non-immunogenic, fixable, easily detected by immunohistochemistry, and because 10kDa (BDA-10) and 70kDa (BDA-70) BDAs exhibit hydrodynamic radii that approximate bioactive protein growth factors and therapeutic monoclonal antibodies, respectively [62, 63].

BDAs injected in PBS are well documented to diffuse locally throughout CNS neural parenchyma and along perivascular spaces, and are taken up by neurons as well as by microglial and perivascular phagocytic cells in a size dependent manner [64]. A detailed comparison of the biodistributions of BDA-10 and BDA-70 released from PBS, DCH_MO_, or DCH_K_ is presented in Supplementary Figures 14-16. Briefly, under all conditions, BDAs were detectable only in the ipsilateral hemisphere, and DCH_MO_, and DCH_K_ displayed significant differences both in the depth of penetration of BDAs into neural tissue and in cellular uptake. BDAs released from DCH_MO_ were found more concentrated locally near hydrogel deposits, with substantial uptake by neurons and some microglial phagocytosis (Supplementary Figs. 14-16). By contrast, BDAs released from DCH_K_ showed increased radial diffusion, but with limited detectable uptake by neurons and heavy accumulation in microglia and perivascular phagocytes (Supplementary Figs. 14-16). These findings suggested that hydrogels with different FBRs exhibit differences not only in the distribution of molecular delivery into neural tissue, but also in the type of cells that may be primarily targeted.

We therefore next compared the biodistribution of BDA-10 released from different hydrogels that displayed escalating and unique severities of FBR: DCH_MO_, DCH_K_, DCH_MM_ and CHIT (Fig. 8a). We quantified the proportion of BDA present in the non-neural compartment defined as the area of hydrogel deposit and its non-neural surrounding tissue, versus the neural compartment defined as the neural tissue surrounding the deposits containing viable neurons and different neuroglia (Fig. 8b-d, Supplementary Fig. 17). At 2 weeks after injection, DCH_MO_, with the least noxious FBR showed the highest total delivery of BDA into the neural compartment, and yielded greater uptake in neurons, which was highest local to the hydrogel-tissue interface (Fig. 8b-d). DCH_K_ exhibited less neural parenchyma delivery than DCH_MO_ but more than DCH_MM_ and CHIT, and BDA was preferentially taken up by microglia rather than neurons (Fig. 8e,f). DCH_MM_ and CHIT, which show the most severe FBR (Fig. 3b,d), displayed: (i) significantly increased accumulation of BDA in non-neural compartments associated with the FBR (Fig. 8b); (ii) reduced uptake of BDA by local neurons (Fig. 8c); and (iii) greater BDA accumulation in macrophages (CD13 positive/CD68 positive cells) (Fig. 8e). Notably, DCH_K_, DCH_MM_ and CHIT exhibited pronounced BBB leakage with extended distribution of serum albumin through neural parenchyma compared with DCH_MO_ (Supplementary Fig. 17c,d) and degree of BBB leak correlated with increased BDA biodistribution throughout the brain and increased inflammatory cell phagocytosis of BDA (Fig. 8d,e, Supplementary Fig. 17a). Serum proteins such as albumin bind systemically delivered biological materials such as bioactive proteins, small molecule drugs and nanoparticles, and alter their biodistribution [65]. In addition, serum proteins are proinflammatory in neural tissue [19, 22]. We also evaluated the biodistribution of BDA-10 released from MP that had been loaded into DCH_MO_. The increased non-neural and phagocyte dominated FBR associated with MP inclusion into nonionic DCH_MO_ correlated with significantly decreased BDA-10 accumulation in neural tissue at 1 week and extensive consumption of BDA-10 by CD13 positive cells (Supplementary Fig. 18). Together, these data show that hydrogels with pronounced non-neural FBRs have reduced molecular delivery efficacy to neural tissue and that this reduction scales with the severity of the FBR.

Lastly, we evaluated the effect of FBR on hydrogel-mediated delivery of bioactive molecules *in vivo*. Basal forebrain cholinergic neurons are exquisitely sensitive to nerve growth factor (NGF) levels and atrophy when deprived of NGF and hypertrophy when exposed to exogenous NGF (Fig. 8g) [34, 66, 67]. To compare the efficacy of NGF delivery by various hydrogels, we quantified the size of local striatal cholinergic neurons. NGF delivered from DCH_MO_ and DCH_K_ stimulated significant 40% increases in ipsilateral cholinergic neuron size relative to uninjected controls, whereas NGF delivered from DCH_MM_ and CHIT, which attract a severe non-neural FBR, showed highly variable effects with no overall significant increase (Fig. 8h). Incorporating data from across the molecular delivery and FBR phenotype characterizations into a Principal Component Analysis (PCA) showed a significant inverse correlation between neural tissue molecular delivery efficacy and the severity and intensity of hydrogel FBR-associated inflammation (Fig. 8i,j, Supplementary Fig. 19).

These findings show that delivery of molecular cargo from hydrogels to CNS tissue is influenced by the nature and severity of the material-evoked FBR (Fig. 9). In particular, an increased recruitment of peripherally derived phagocytes into hydrogel deposits results in increased consumption of cargo molecules and reduced neural tissue delivery. Further, a minimal hydrogel FBR with little or no BBB leakage results in very local delivery with a higher proportion of targeting of cargo molecules to neurons. In contrast, more severe FBRs with pronounced BBB leakage contributes to increased dispersal of delivered molecules throughout larger volumes of neural parenchyma, and to greater targeting of those molecules to phagocytosis by microglia and perivascular cells.

## Discussion

Here, we show that CNS FBRs to biomaterials mimic CNS wound responses and exist on a severity spectrum that is determined by definable chemical functionalities perceived by the host at the biomaterial-tissue interface. Moreover, the nature and severity of FBRs significantly affect multiple aspects of biomaterial function. These findings have broad implications for developing, testing and using biomaterials for CNS applications, and also for understanding the conserved biology of the CNS wound response and astroglial limitans border formation. We provide a framework for comprehensively evaluating *in vivo* responses to biomaterials for CNS applications, which may facilitate the development and characterization of new and improved systems.

As a platform to study CNS FBRs evoked by definable chemical functionalities presented to host cells by biomaterials, we used readily modified synthetic hydrogels whose direct interfaces with host cells can sensitively and intimately be evaluated *in vivo*. Our findings show that specific physiochemical properties presented at biomaterial-host interfaces, such as cationic, anionic or nonionic interfaces with CNS cells, evoke quantifiably different FBR phenotypes with respect to levels of stromal cell infiltration, inflammation, tissue fibrosis, astroglial border formation and BBB disruption. Moreover, we show that FBRs evoked by materials in CNS are directly related to, and mimic, the multicellular CNS wound response to tissue injury (modeled here by the response to an ischemic infarct), and are best understood in the context of this conserved biology. Trauma, ischemia, autoimmune attack and infections all generate potentially toxic cellular debris that trigger multicellular wound responses as naturally occurring events. The conserved CNS wound response serves to recruit inflammatory cells that degrade toxic elements and also simultaneously serves to isolate damaged tissue, debris and inflammation from adjacent viable neural tissue and to restore barrier functions important for normal CNS function [19, 22, 55]. In this manner, naturally occurring CNS wound responses give rise to discrete lesion compartments with central cores of non-neural tissue surrounded by astrocyte limitans borders that are continuous with spared and reorganizing neural tissue. This compartmentalized organization is mimicked by FBRs evoked by biomaterials (Fig. 9). The specialized multicellular and compartmentalized CNS wound response or FBR is not present in peripheral tissues, and thus evaluation of biomaterial FBR in peripheral tissue alone is not sufficient to predict performance in the CNS. Naturally occurring CNS wound responses exist on a severity spectrum that varies with: (i) the degree of the tissue damage, (ii) the nature and the toxicity of debris, and (iii) the presence or absence of microbes or their associated signals. Similarly, the nature and severity of FBRs to biomaterials vary with the molecular features of the interface presented to host tissue.

Our findings emphasize the need for use of multiple markers to assess multiple FBR dimensions, including markers of stromal, inflammatory, multiple glial cells, neural tissue damage and fibrosis. We show that GFAP, a commonly used marker of astrocyte reactivity that can indicate tissue pathology [19], is on its own not a sufficient or reliable marker to evaluate and differentiate material or device FBRs. For example, we show that total GFAP levels are essentially indistinguishable in neural tissue within close proximity to biomaterials that evoke markedly different FBR severities, including FBRs that evoke pronounced levels of inflammation and fibrosis that impact significantly on function. Our findings provide a framework of multiple molecular markers and experimental tools for comprehensive evaluation of material FBRs in the CNS *in vivo* using readily available immunohistochemical reagents that can be applied to benchmark the performance of new biomaterial formulations against various well characterized formulations used here.

We also show that the nature and severity of FBRs strongly influence biomaterial function in the CNS as studied here in the form of rate of hydrogel resorption, biodistribution of molecular cargo delivery to neural parenchyma and efficacy of bioactive molecular delivery. Thus, developing hydrogels and other biomaterials with efficient functions for CNS applications will require a detailed understanding of their *in vivo* FBRs. *In vitro* evaluations of molecular release dynamics are on their own insufficient to predict efficacy of molecular delivery in the CNS *in vivo*, because delivery will be strongly influenced by CNS specific FBRs. For example, we show that microparticles that significantly extend molecular release duration from hydrogels *in vitro* can attract phagocytes *in vivo* that significantly reduce molecular delivery compared with the hydrogels they are suspended in. Notably, our findings indicate that hydrogel formulations with features that attract phagocytic cells such as blood borne macrophages and neutrophils will have particularly poor functional performance in molecular delivery to host tissue (and most likely also as vehicles for cell grafts). Moreover, we show that these hydrogel features may not be revealed by solely examining the glial response of more distant surrounding neural tissue, because hydrogel deposits are rapidly and effectively isolated as part of the FBR, which mimics naturally occurring wound responses that maximize the protection of adjacent neural tissue [19, 22].

In this study we used hydrogels as surrogate biomaterials to characterize host responses to specifically presented chemical functionalities, and molecular delivery from hydrogels was the primary functional outcome measure evaluated. The basic principles we identified can inform the development of materials for a variety of CNS applications including implantable neuroprostheses for recording and stimulating neural circuit activity [2, 3, 23, 68]. For example, we show that surface charge at the material-host interface influences the severity of FBRs, which in turn will influence the degree of protective fibrotic and astrocyte barrier formation that forms to isolate the materials from adjacent neural tissue and may thereby attenuate function. Thus, such devices will also benefit from detailed characterization of FBRs as described here. While the bulk of implantable neuroprostheses most frequently employed in human and non-human primate studies currently make use of non-resorbable, hydrophobic polymers that may stimulate unique FBR phenotypes in addition to that described here [23], emerging systems are opting for hydrophilic, hydrogel-based coatings/substrates with comparable chemistries to this study [69, 70]. Thus, it would seem particularly appropriate to apply the FBR evaluation framework outlined here to these new formulations as part of preclinical testing protocols moving forward. Furthermore, incorporating the identified hydrogel chemical functionalities that evoked minimal FBRs (e.g. methionine sulfoxide functional groups from DCH_MO_) into future neural electrode design may be a useful strategy in attenuating the thickness of disrupted tissue and the intensity of the astroglial border-forming response between the device and viable neural tissue.

Lastly, our findings inform the understanding of CNS wound responses by demonstrating and characterizing how FBRs fall firmly within the framework of the conserved multicellular biology of such responses. In this regard, we also add to the growing evidence [48–53] that APP accumulation and the acute formation of Aβ are consequent responses to neural tissue injury that occur in direct proportion to the level of neuronal and axonal damage, and are thus part of the conserved biology of CNS wound responses and FBRs.

In conclusion, we show that material FBRs are determined by definable properties presented at host interfaces, which can be modified to minimize or evoke specific responses. Understanding these properties and the CNS specific FBRs that they evoke *in vivo* will be critical to designing new materials for diverse CNS applications. In addition, this observation raises the intriguing prospect of engineering biomaterials with originally inert profiles to present specific molecular features in order to dissect cell-surface and contact-mediated molecular mechanisms that drive specific elements of naturally occurring CNS wound responses.

## Methods

### Animals

All *in vivo* experiments involving the use of mice were conducted according to protocols approved by the Animal Research Committee (ARC) of the Office for Protection of Research Subjects at University of California Los Angeles (UCLA). All *in vivo* animal experiments were conducted within approved UCLA facilities using wildtype or transgenic C57/BL6 female and male mice that were aged between 8 weeks and four months old at the time of craniotomy surgery. Transgenic td-Tomato (tdT) reporter mice were bred with inducible Cre lines to identify Astrocytes (Aldh1L1-Cre-ERT2) and NG2 cells (NG2-Cre-ERT2). Transgenic mice (aged between 6 and 12 weeks) were administered tamoxifen to induce Cre expression. Tamoxifen in corn oil at a concentration of 20mg/ml was administered via intraperitoneal (IP) injection for 5 days at a dose of 50mg/kg per day. Following the last dose of tamoxifen, mice were kept for 3 weeks before undergoing surgery for hydrogel injection to allow residue tamoxifen to clear out of the mouse’s system. Mice were housed in a 12-hour light/dark cycle in a specific pathogen-free facility with controlled temperature and humidity and were provided with food and water ad libitum.

### Surgical procedures for mice

All surgical procedures were approved by the UCLA ARC and conducted within a designated surgical facility at UCLA. Hydrogel brain injections were performed on mice under general anesthesia achieved through inhalation of isoflurane in oxygen-enriched air. Prior to incision, the mouse head was stabilized and horizontally leveled in a rodent stereotaxic apparatus using ear bars (David Kopf, Tujunga, CA). A small craniotomy was performed using a high speed surgical drill and visually aided by an operating microscope (Zeiss, Oberkochen, Germany). Hydrogel injections were made in the brain using pulled borosilicate glass micropipettes (WPI, Sarasota, FL, #1B100-4) that were ground to a 35° beveled tip with 150–250 μm inner diameter. Glass micropipettes were mounted to the stereotaxic frame via specialized connectors and attached, via high-pressure polyetheretherketone (PEEK) tubing, to a 10 μL syringe (Hamilton, Reno, NV, #801 RN) that was controlled by an automated syringe pump (Pump 11 Elite, Harvard Apparatus, Holliston, MA). Each hydrogel was backloaded into the micropipette prior to connecting to the stereotaxic frame. A volume of 1 μL of hydrogel was injected into the caudate putamen nucleus at 0.15 μL/min using target coordinates relative to Bregma: +0.5 mm A/P, +2.5 mm L/M and −3.0 mm D/V. For the NGF delivery studies target coordinates relative to Bregma: +1 mm A/P, +2.2 mm L/M and −3.0 mm D/V were used. The micropipette was allowed to dwell in the brain at the injection site for an additional 4 minutes at completion of hydrogel injection. The micropipette was then removed from the brain slowly and incrementally over a 2 minute period.

### Synthesis and formulation of hydrogels

Physically crosslinked hydrogels were formulated from polypeptide or polysaccharide polymers (Figure 1a, Supplementary Fig. 3 a,b) that were synthesized using procedures developed previously by our group or purchased from commercial sources and used as received.

#### Preparation of Diblock Copolypeptide Hydrogels (DCH)

The polypeptides were prepared using synthetic schemes reported previously. Specific and detailed synthetic protocols for preparing DCH_MO_, DCH_K_, DCH_MM_, DCH_E_, DCH_MOK10_, DCH_MOE15_ have been provided across a number of previous peer reviewed publications [8, 25, 34].

DCH_MOE15_ was prepared for the first time as part of this current work and the method for synthesizing this polypeptide can serve as a general overview for polypeptide synthesis. Polypeptide synthesis was performed in a N_2_ filled glove box using anhydrous solvents. Amino acid N-carboxyanhydride (NCA) monomers including *tert*-butyl-L-glutamate NCA, L-methionine NCA, L-leucine NCA were prepared by phosgenation of a tetrahydrofuran (THF) solution of the corresponding amino acid in a Schlenk flask under N_2_ and purified by either recrystallization or column chromatography as outlined before. To prepare copolypeptides at ca. 100 mg scale, a solution of initiator, Co(PMe_3_)_4_, (3.5 mg, 0.001 mmol) in THF (20 mg/mL) was quickly added to a mixture of L-methionine NCA (Met NCA; 100 mg, 0.57 mmol) and *tert*-butyl-L-glutamate NCA (*tert*-butyl Glu NCA; 23 mg, 0.10 mmol) in THF (50 mg/mL). After 2 hours, complete consumption of NCA was confirmed by Fourier-transform infrared (FTIR) spectroscopy. In order to determine the lengths of poly(L-methionine-*stat*-*tert*-butyl-L-glutamate)_m_, (ME^tB^)m, segments, a small aliquot (200 µl) of polymerization mixture was removed, reacted with α-methoxy-ω-isocyanoethyl-poly(ethylene glycol)_45_ (mPEG_23_-NCO) and chain length determined by end-group analysis using ^1^H NMR. To the remaining copolymerization mixture L-leucine NCA (Leu NCA; 14 mg, 0.088 mmol) in THF (50 mg/mL) was then added. The Leu NCA monomers were found to be completely consumed within approximately 1 hour, and the reaction mixture was subsequently removed from the glove box. The block copolypeptide solution was then precipitated by addition to a DI water solution (75 mL), filtered, washed with DI water, and dried under reduced pressure to obtain a white fluffy solid (91 mg, 95% yield).

The deprotection of *tert*-butyl-L-glutamate residues was performed before the oxidation of L-methionine residues. To facilitate deprotection, (ME^tB^)_170_L_24_ (ca. 80 mg) was fully dissolved in trifluoroacetic acid (*ca*. 25 mg/mL) and stirred at ambient temperature for 16 hours. The copolypeptide was then precipitated into diethyl ether (75 mL). The diethyl ether was decanted and the copolypeptide was dried under reduced pressure overnight to yield a white solid (*ca*. 74 g, 99% yield, 97% deprotection efficiency). To convert L-methionine residues in copolypeptides to L-methionine sulfoxide residues, a volume of 70 wt. % *tert*-butyl hydroperoxide (TBHP) (16 molar equivalents per L-methionine residue) was added to a sample of solid copolypeptide (ca. 80 mg). The reaction mixture was then diluted with DI water to yield an overall polypeptide concentration of 20 mg/mL. To aid oxidation, a catalytic amount of camphorsulfonic acid (CSA) (0.2 molar equivalents per Met residue) solution in DI water (20 mg/ml) was subsequently added. The reaction mixture was stirred vigorously for 24 hours at ambient temperature, whereupon complete dissolution of the copolypeptide sample was observed. Reaction mixtures were transferred to 2000 MWCO dialysis bags and dialyzed against: (i) pyrogen free deionized milli-Q water (3.5 L) containing sodium thiosulfate (1.2 g, 2.2 mM) for 1 day to neutralize residual peroxide, (ii) pyrogen free deionized milli-Q water (3.5 L) containing ethylenediaminetetraacetic acid tetrasodium salt hydrate (1.0 g, 2.63 mmol) to aid cobalt ion removal, (iii) pyrogen free deionized milli-Q water (3.5 L) which was adjusted to pH 8 with 1.0 M NaOH and contained sodium chloride (5 g, 85.5 mmol) for 2 days to aid in counter ion exchange of glutamate residues, and (IV) pyrogen free deionized milli-Q water (3.5 L) for 2 days to remove residual ions. For each step dialysate was changed every 12 hours. The copolypeptide solution was then freeze dried to yield a white fluffy solid (ca. 85 g, 95% yield, and 100% conversion of L-methionine residues in copolypeptides to L-methionine sulfoxide groups).

All DCH variants were prepared by solubilizing lyophilized copolypeptide in phosphate buffered saline (PBS) and leaving to assemble for 24 hours, without stirring, before use. All DCH formulations were prepared at 4.0 wt% in PBS.

#### Preparation of Polysaccharide based hydrogels

Polysaccharide based hydrogels of chitosan, methyl cellulose (MC) and hyaluronic acid/methyl cellulose blend (HAMC) were prepared following procedures detailed in the literature. To formulate chitosan hydrogels, solutions of chitosan [MW= 50,000-190,000 Da; DDA≈85%] (Sigma, #448869) in 0.1M acetic acid were prepared at 2.3 % (w/v) and allowed to dissolve overnight at 4°C [33]. A 50% w/v or 2.314 M solution of β-glycerophosphate disodium salt (β-GP) (Sigma, #G9422) in water was subsequently added dropwise to the chitosan solution on ice and manually stirred to form a uniform solution and elevate the solution pH to 7.4. Prior to surgical micropipette loading, the chitosan hydrogel was incubated at 37°C for one hour to initiate physical gelation. Methylcellulose (Methocel A15C, [MW = 304 kDa; 27.5-31.5% substitution], Sigma #64625) hydrogels were prepared by dispersing and then solubilizing at 4°C at a concentration of 4 wt% [31]. The HAMC hydrogel was prepared by dispersing a measured amount of hyaluronic acid [1.01-1.8 MDa] (Lifecore Biomedical, #HA15M-1) into a pre-dissolved 4wt% MC solution and allowed to dissolve at 4°C overnight. The HAMC blended hydrogel used for *in vivo* studies had a concentration of 1% HA and 3% MC which was equivalent that used previously within *in vivo* CNS studies [32].

For BDA delivery experiments, appropriate volumes of 10% w/v stock solutions of BDA (10kDa – Thermofisher, #D1956 or 70kDa – Thermofisher, # D1957) in PBS were added to the polymer solutions to net a final concentration of 1% BDA in hydrogels. For NGF delivery experiments, lyophilized recombinant human β-NGF (PeproTech, Cat# 450-01, Rocky Hill, NJ) was reconstituted in PBS and added to the polymers to form hydrogels with a final concentration of NGF of 1 mg/mL.

### Microgel Microparticle Synthesis

Microparticles based on polyethylene glycol (PEG) microgels were derived by covalently reacting ethoxylated trimethylolpropane (TMPE) oligomers functionalized with either acrylate or thiol acid ester groups under physiologically buffered conditions by thiol-ene Michael Addition as reported before [58]. A standard inverse emulsion (IE) technique was applied to formulate microgels whereby aqueous buffer solubilized oligomers were dispersed in a larger volume mineral oil bath that contained the nonionic sorbitan alkyl ester surfactant, Span 83, to create a stable emulsion and direct microgel size [59, 60]. The emulsion was stirred for one hour at room temperature on a magnetic stir plate before being washed and centrifuged three times with hexanes to remove the mineral oil and surfactant. Collected microparticles were dried under high vacuum to evaporate residual hexane solvent. Microgels were uniformly dispersed in PBS at 6 wt% using a pellet pestle mixer and added to hydrogel solutions to a final concentration of 3 wt%. For BDA delivery experiments, appropriate volumes of 10% BDA-10 stock were added to the aqueous reaction buffer during microgel synthesis to load microgel particles with BDA.

### Hydrogel and Microparticle Characterization

#### Dynamic Mechanical Rheology

Dynamic rheological measurements were performed at constant set temperature of 25 °C using a Physica MCR 301 (Anton Paar, Sweden) equipped with a 25 mm 1° cone and plate geometry. To determine the linear viscoelastic testing region for each DCH sample a frequency sweep from 0.1 to 100 rad s^−1^ at a fixed strain of 0.5% was applied followed by a strain sweep from 0.1 to 100% strain at a fixed frequency of 10 rad s^−1^. DCH samples were allowed to recover for a period of 2 minutes between individual rheological experiments. To compare mechanical properties across various DCH samples a time sweep under fixed strain and frequency conditions (0.5% strain and 10 rad s^−1^) was applied for one minute.

#### Zeta potential Measurement

K_170_ (hydrophilic block of DCH_K_), (M^O^A)_170_ (hydrophilic block of DCH_MO_), and (M^M^A)_170_ (hydrophilic block of DCH_MM_), were dissolved at 15 mg/mL in a 20 mM phosphate buffer solution at pH 7.4 which was filtered through a 0.45 µm membrane syringe filter. Mean zeta potentials were analyzed using a Zetasizer NanoZS instrument (Malvern Instruments Ltd., United Kingdom). Samples were analyzed in monomodal mode where fast field reversal (FFR) technique was applied and the electrophoretic mobility was calculated using the Henry equation. Sample were prepared in triplicate and analyzed 3 times.

#### Fourier Transform Infrared Spectroscopy (FTIR)

Dried microparticles were analyzed by FTIR to confirm the presence of the crosslinked ethoxylated polyol network. Several milligrams of dried microparticles were analyzed by attenuated total reflection Fourier transform infrared (ATR-FTIR) spectroscopy using a PerkinElmer Spectrum One instrument equipped with a universal ATR assembly. The ester stretch (≈1730 cm^−1^) and the loss of the acrylate C=C stretch (1675–1600 cm^−1^) were used to confirm presence of the covalent crosslinked network [58].

#### Differential interference contrast (DIC) microscopy

To visualize and quantify the size of formulated microparticles a 1mg/ml solution of dispersed microparticles was placed on a glass slide, coverslipped and imaged on a Zeiss Axioplan photomicroscope using a 63X oil objective. Quantification of microparticles size was performed on images taken from several separately prepared slides using NIH Image J (1.51) software.

### Immunohistochemistry

After terminal anesthesia by barbiturate overdose mice were perfused transcardially with heparinized saline and 4% paraformaldehyde. Brains were immediately dissected after perfusion and post-fixed in 4% PFA for 6 hours. Brains were cryoprotected in 30% sucrose in TBS for at least 3 days with the sucrose solution replaced once after 2 days. Coronal brain sections (40 µm thick) were cut using a Leica CM3050 cryostat. Brain sections were stored transiently in TBS buffer at 4°C or in antifreeze solution of 50% glycerol/30% Sucrose in PBS at -20°C for long term storage. Brain sections were processed for immunofluorescence as described previously [4, 5]. The primary antibodies used in this study were: rabbit anti-GFAP (1:1000; Dako, Santa Clara, CA); rat anti-GFAP (1:1000, Thermofisher, Grand Island, NY); rabbit anti NeuN (1:1000, Abcam, Cambridge, MA); guinea pig anti-NeuN (1:1000, Synaptic Systems, Goettingen, Germany); goat anti-CD13 (1:200, R&Dsystems, Minneapolis, MN); rabbit anti-Laminin 1 (1:100, Sigma, St.Louis, MO); rabbit anti-Fibronectin (1:500, Millipore, Burlington, MA); rabbit anti-Collagen 1a1 (1:300, Novus Biologicals, Littleton, CO); rabbit anti-RFP (1:1000, Rockland, Limerick, PA); goat anti-Albumin (1:300, Novus Biologicals, Littleton, CO); rat anti-PECAM-1 (1:200, BD Biosciences, San Jose, CA); rat anti-Galectin-3 (1:200, Invitrogen-Thermofisher Scientific, Grand Island, NY); rat anti-CD68 (1:1000, AbDserotec-BioRad, Hercules, CA); rat anti-CD45 (1:100, BD Biosciences, San Jose, CA); rabbit anti-Iba-1 (1:800, Wako, Osaka, Japan); guinea pig anti-Iba-1 (1:800, Synaptic systems, Goettingen, Germany); Rabbit anti-P2Y12R (1:500, Anaspec, Fremont, CA); rabbit anti-mouse IgG (1:1000, Abcam, Cambridge, MA); rat anti-Ly6B2 (1:200, Bio-Rad, Hercules, CA); Goat anti-PDGFR-β (1:200, R&Dsystems, Minneapolis, MN); and rabbit anti-Olig2 (1:200, Millipore, Burlington, MA). BDA was visualized with streptavidin-HRP plus Tyr-Cy3 (Perkin Elmer) as before [4]. Nuclei were stained with 4’,6’-diamidino-2-phenylindole dihydrochloride (DAPI; 2ng/ml; Molecular Probes). Sections were coverslipped using ProLong Gold anti-fade reagent (InVitrogen, Grand Island, NY). Sections were examined and photographed using epifluorescence microscopy, deconvolution fluorescence microscopy and scanning confocal laser microscopy (Zeiss, Oberkochen, Germany). Tiled scans of individual whole sections were prepared using a x20 objective and the scanning function of a Leica Aperio Versa 200 Microscope (Leica, Wetzlar, Germany) available in the UCLA Translational Pathology Core Laboratory.

### Quantification of Immunohistochemistry

Immunohistochemical staining intensity quantification was performed on whole brain images derived from the slide scanner or tiled images prepared on the epifluorescence microscope. All images used for each comparable analysis were taken at a standardized exposure time and used the raw/uncorrected intensity setting. Quantification of antibody staining intensity for the individual hydrogels was performed using NIH Image J (1.51) software and the Radial Profile Angle plugin. The procedure for determining the hydrogel radius and the non-neural and neural compartments is outlined graphically in Supplementary Fig. 2. Briefly, a radial profile field of 900µm (~846 pixels) was applied to each hydrogel (this standardized radial profile field was determined as the average field that included all hydrogels and was confined within the caudate putamen region of the brain section (i.e. between the ventricle and corpus callosum)). The radial profile analysis determines the average pixel intensity value across the circumference of a circle at each radial pixel distance. The Hydrogel radius/start of tissue location was determined from the DAPI profile and specifically where the first derivative (dI/dx) of the DAPI profile crossed a threshold of +0.2. The non-neural/neural tissue boundary was defined as the maximum of the GFAP intensity which was determined where dI/dx equaled zero. Total values for CD13, GFAP, Ly6B2, Col1a1, Gal-3, Albumin, IgG, Iba-1, BDA stainings were determined by taking the integral (Area under the curve (AUC)) of the radial intensity profile. Albumin and IgG intensities and total values were normalized to the contralateral hemisphere of each animal.

For BDA and NGF delivery quantification only slide scanned images were used. Medial and lateral hem-circular radial analysis was performed in addition to the standard radial profile analysis described above to quantify BDA dispersion throughout whole brain sections. Quantification of the co-staining of BDA with specific immunohistochemical stains such as NeuN, CD13, CD68, Iba-1, IgG and Albumin was performed using the RG2B colocalization plugin in NIH Image J (1.51) software. Number of BDA and NeuN positive neurons was determined by counting the number of BDA/NeuN colocalized staining puncta using the analyze particles function in NIH Image J (1.51) software. Bioactive NGF delivery was assessed by characterizing the size (area) of Choline Acetyltransferase (ChAT) and NeuN positive cholinergic neurons also using the analyze particles function in NIH Image J (1.51) software.

### Statistics, power calculations, group sizes and reproducibility

Graph generation and statistical evaluations of repeated measures were conducted by one-way or two-way ANOVA with post hoc independent pair wise analysis as per Bonferroni, Tukey, or by Student’s t-tests where appropriate using Prism 8 (GraphPad Software Inc, San Diego, CA). Principal Component Analysis (PCA) was performed using XLStat (Addinsoft Inc, Long Island City, NY). For one-way and two-way ANOVA statistical evaluations, P-values, F-values and degrees of freedom are reported in the manuscript and within the source data files. Similarly, for Student’s two-tailed t-test, the t value and degrees of freedom are reported. Power calculations were performed using G*Power Software V 3.1.9.2. For immunohistochemical quantification analysis, group sizes were calculated to provide at least 80% power when using the following parameters: probability of type I error (alpha) = .05, a conservative effect size of 0.25, 3-10 treatment groups with multiple measurements obtained per replicate. All graphs show mean values plus or minus standard error of the means (S.E.M.) as well as individual values as dot plots. All bar graphs are overlaid with dot plots where each dot represents the value for one animal to show the distribution of data and the number (n) of animals per group. Files of all individual values are provided as source data. Injections of hydrogel formulations were repeated independently at least three times in different colonies of mice across a two-year period with similar results.

### Data availability statement

Files of source data of individual values for all quantitative figures are provided with the paper. Other data that support the findings of this study are available on reasonable request from the corresponding author.

## Acknowledgements

This work was supported by US National Institutes of Health (NS084030 to M.V.S.); Dr. Miriam and Sheldon G. Adelson Medical Foundation (M.V.S. and T.J.D); Craig H. Neilsen Foundation (381357 to T.M.O.); Paralyzed Veterans Foundation of America (RF3170 to T.M.O.); American Australian Association (T.M.O.), Wings for Life Spinal Cord Research Foundation (M.V.S. and T.M.O.); and Microscopy Core Resource of UCLA Broad Stem Cell Research Center.

## Author contributions

T.M.O., T.J.D. and M.V.S. designed experiments; T.M.O., A.L.W. J.H.K. and Y.A. conducted experiments; T.M.O., A.L.W, T.J.D. and M.V.S. analyzed data. T.M.O., T.J.D. and M.V.S. prepared the manuscript.

## Statement about Competing interests

The authors declare no competing interests.

## References

1. Orive, G., et al., Biomaterials for promoting brain protection, repair and regeneration. Nature Reviews Neuroscience, 2009. 10: p. 682.

2. Zhong, Y. and R.V. Bellamkonda, Biomaterials for the central nervous system. Journal of the Royal Society, Interface, 2008. 5(26): p. 957–975.

3. Obidin, N., F. Tasnim, and C. Dagdeviren, The Future of Neuroimplantable Devices: A Materials Science and Regulatory Perspective. Advanced Materials. 0(0): p. 1901482.

4. Anderson, M.A., et al., Required growth facilitators propel axon regeneration across complete spinal cord injury. Nature, 2018. 561(7723): p. 396–400.

5. Anderson, M.A., et al., Astrocyte scar formation aids central nervous system axon regeneration. Nature, 2016. 532: p. 195.

6. O’Shea, T.M., et al., CHAPTER 19 Smart Materials for Central Nervous System Cell Delivery and Tissue Engineering, in Smart Materials for Tissue Engineering: Applications. 2017, The Royal Society of Chemistry. p. 529–557.

7. Mitrousis, N., A. Fokina, and M.S. Shoichet, Biomaterials for cell transplantation. Nature Reviews Materials, 2018. 3(11): p. 441–456.

8. Wollenberg, A.L., et al., Injectable polypeptide hydrogels via methionine modification for neural stem cell delivery. Biomaterials, 2018. 178: p. 527–545.

9. Slaughter, B.V., et al., Hydrogels in regenerative medicine. Advanced materials (Deerfield Beach, Fla.), 2009. 21(32-33): p. 3307–3329.

10. Lacour, S.P., G. Courtine, and J. Guck, Materials and technologies for soft implantable neuroprostheses. Nature Reviews Materials, 2016. 1: p. 16063.

11. Kozai, T.D.Y., et al., Brain Tissue Responses to Neural Implants Impact Signal Sensitivity and Intervention Strategies. ACS Chemical Neuroscience, 2015. 6(1): p. 48–67.

12. Green, R.A., et al., Conducting polymers for neural interfaces: Challenges in developing an effective long-term implant. Biomaterials, 2008. 29(24): p. 3393–3399.

13. Elliott Donaghue, I., et al., Cell and biomolecule delivery for tissue repair and regeneration in the central nervous system. J Control Release, 2014. 190: p. 219–27.

14. Kearney, C.J. and D.J. Mooney, Macroscale delivery systems for molecular and cellular payloads. Nature Materials, 2013. 12: p. 1004.

15. Loebel, C., et al., Shear-thinning and self-healing hydrogels as injectable therapeutics and for 3D-printing. Nature Protocols, 2017. 12: p. 1521.

16. Wang, L.L., et al., Sustained miRNA delivery from an injectable hydrogel promotes cardiomyocyte proliferation and functional regeneration after ischaemic injury. Nature Biomedical Engineering, 2017. 1(12): p. 983–992.

17. Zhang, L., et al., Zwitterionic hydrogels implanted in mice resist the foreign-body reaction. Nature Biotechnology, 2013. 31: p. 553.

18. Appel, E.A., et al., Self-assembled hydrogels utilizing polymer–nanoparticle interactions. Nature Communications, 2015. 6: p. 6295.

19. Burda, J.E. and M.V. Sofroniew, Reactive gliosis and the multicellular response to CNS damage and disease. Neuron, 2014. 81(2): p. 229–48.

20. Bellver-Landete, V., et al., Microglia are an essential component of the neuroprotective scar that forms after spinal cord injury. Nature Communications, 2019. 10(1): p. 518.

21. Brennan, F.H., et al., Complement receptor C3aR1 controls neutrophil mobilization following spinal cord injury through physiological antagonism of CXCR2. JCI Insight, 2019. 4(9).

22. O’Shea, T.M., J.E. Burda, and M.V. Sofroniew, Cell biology of spinal cord injury and repair. The Journal of clinical investigation, 2017. 127(9): p. 3259–3270.

23. Salatino, J.W., et al., Glial responses to implanted electrodes in the brain. Nature biomedical engineering, 2017. 1(11): p. 862–877.

24. Nowak, A.P., et al., Rapidly recovering hydrogel scaffolds from self-assembling diblock copolypeptide amphiphiles. Nature, 2002. 417(6887): p. 424–8.

25. Yang, C.-Y., et al., Biocompatibility of amphiphilic diblock copolypeptide hydrogels in the central nervous system. Biomaterials, 2009. 30(15): p. 2881–2898.

26. Webber, M.J., et al., Supramolecular biomaterials. Nature Materials, 2015. 15: p. 13.

27. Nunez, S., et al., A Versatile Murine Model of Subcortical White Matter Stroke for the Study of Axonal Degeneration and White Matter Neurobiology. Journal of visualized experiments : JoVE, 2016(109): p. 53404.

28. Shipp, M.A. and A.T. Look, Hematopoietic differentiation antigens that are membrane-associated enzymes: cutting is the key! Blood, 1993. 82(4): p. 1052–70.

29. Armulik, A., et al., Pericytes regulate the blood–brain barrier. Nature, 2010. 468: p. 557.

30. Bush, T.G., et al., Leukocyte Infiltration, Neuronal Degeneration, and Neurite Outgrowth after Ablation of Scar-Forming, Reactive Astrocytes in Adult Transgenic Mice. Neuron, 1999. 23(2): p. 297–308.

31. Tate, M.C., et al., Biocompatibility of methylcellulose-based constructs designed for intracerebral gelation following experimental traumatic brain injury. Biomaterials, 2001. 22(10): p. 1113–1123.

32. Gupta, D., C.H. Tator, and M.S. Shoichet, Fast-gelling injectable blend of hyaluronan and methylcellulose for intrathecal, localized delivery to the injured spinal cord. Biomaterials, 2006. 27(11): p. 2370–9.

33. Crompton, K.E., et al., Inflammatory response on injection of chitosan/GP to the brain. Journal of Materials Science: Materials in Medicine, 2006. 17(7): p. 633–639.

34. Song, B., et al., Sustained local delivery of bioactive nerve growth factor in the central nervous system via tunable diblock copolypeptide hydrogel depots. Biomaterials, 2012. 33(35): p. 9105–16.

35. Jeong, H.-K., et al., Inflammatory Responses Are Not Sufficient to Cause Delayed Neuronal Death in ATP-Induced Acute Brain Injury. PLOS ONE, 2010. 5(10): p. e13756.

36. Sasaki, Y., et al., Selective expression of Gi/o-coupled ATP receptor P2Y12 in microglia in rat brain. Glia, 2003. 44(3): p. 242–250.

37. Haynes, S.E., et al., The P2Y12 receptor regulates microglial activation by extracellular nucleotides. Nature Neuroscience, 2006. 9(12): p. 1512–1519.

38. Kendall, R.T. and C.A. Feghali-Bostwick, Fibroblasts in fibrosis: novel roles and mediators. Frontiers in Pharmacology, 2014. 5(123).

39. Fernandez-Klett, F., et al., Early loss of pericytes and perivascular stromal cell-induced scar formation after stroke. J Cereb Blood Flow Metab, 2013. 33(3): p. 428–39.

40. Aldrich, A. and T. Kielian, Central nervous system fibrosis is associated with fibrocyte-like infiltrates. Am J Pathol, 2011. 179(6): p. 2952–62.

41. Henderson, N.C., et al., Galectin-3 regulates myofibroblast activation and hepatic fibrosis. Proc Natl Acad Sci U S A, 2006. 103(13): p. 5060–5.

42. Lalancette-Hébert, M., et al., Galectin-3 Is Required for Resident Microglia Activation and Proliferation in Response to Ischemic Injury. The Journal of Neuroscience, 2012. 32(30): p. 10383–10395.

43. Fernández-Klett, F. and J. Priller, The Fibrotic Scar in Neurological Disorders. Brain Pathology, 2014. 24(4): p. 404–413.

44. Li, L.-c., J. Li, and J. Gao, Functions of Galectin-3 and Its Role in Fibrotic Diseases. Journal of Pharmacology and Experimental Therapeutics, 2014. 351(2): p. 336–343.

45. Yan, Y.-P., et al., Galectin-3 mediates post-ischemic tissue remodeling. Brain Research, 2009. 1288: p. 116–124.

46. Sirko, S., et al., Astrocyte reactivity after brain injury—: The role of galectins 1 and 3. Glia, 2015. 63(12): p. 2340–2361.

47. Rakers, C., et al., Stroke target identification guided by astrocyte transcriptome analysis. Glia, 2019. 67(4): p. 619–633.

48. Hortobagyi, T., et al., Traumatic axonal damage in the brain can be detected using beta-APP immunohistochemistry within 35 min after head injury to human adults. Neuropathol Appl Neurobiol, 2007. 33(2): p. 226–37.

49. O’Brien, R.J. and P.C. Wong, Amyloid precursor protein processing and Alzheimer’s disease. Annual review of neuroscience, 2011. 34: p. 185–204.

50. Sokrab, T.-E.O., et al., A transient hypertensive opening of the blood-brain barrier can lead to brain damage. Acta Neuropathologica, 1988. 75(6): p. 557–565.

51. Sherriff, F.E., L.R. Bridges, and S. Sivaloganathan, Early detection of axonal injury after human head trauma using immunocytochemistry for beta-amyloid precursor protein. Acta Neuropathol, 1994. 87(1): p. 55–62.

52. Kumar, D.K.V., et al., Amyloid-β peptide protects against microbial infection in mouse and worm models of Alzheimer’s disease. Science translational medicine, 2016. 8(340): p. 340ra72–340ra72.

53. Wu, Y., et al., Microglia and amyloid precursor protein coordinate control of transient Candida cerebritis with memory deficits. Nature Communications, 2019. 10(1): p. 58.

54. Zhu, X., et al., Age-dependent fate and lineage restriction of single NG2 cells. Development, 2011. 138(4): p. 745–53.

55. Sofroniew, M.V., Astrocyte barriers to neurotoxic inflammation. Nature Reviews Neuroscience, 2015. 16: p. 249.

56. Baumann, M.D., et al., An injectable drug delivery platform for sustained combination therapy. J Control Release, 2009. 138(3): p. 205–13.

57. Mura, S., J. Nicolas, and P. Couvreur, Stimuli-responsive nanocarriers for drug delivery. Nature Materials, 2013. 12: p. 991.

58. O’Shea, T.M., et al., Synthesis and Characterization of a Library of In-Situ Curing, Nonswelling Ethoxylated Polyol Thiol-ene Hydrogels for Tailorable Macromolecule Delivery. Advanced Materials, 2015. 27(1): p. 65–72.

59. Murthy, N., et al., A macromolecular delivery vehicle for protein-based vaccines: Acid-degradable protein-loaded microgels. Proceedings of the National Academy of Sciences, 2003. 100(9): p. 4995–5000.

60. Tibbitt, M.W., et al., Student award for outstanding research winner in the Ph.D. category for the 9th World Biomaterials Congress, Chengdu, China, June 1–5, 2012. Journal of Biomedical Materials Research Part A, 2012. 100A(7): p. 1647–1654.

61. Champion, J.A., A. Walker, and S. Mitragotri, Role of Particle Size in Phagocytosis of Polymeric Microspheres. Pharmaceutical Research, 2008. 25(8): p. 1815–1821.

62. Wolak, D.J. and R.G. Thorne, Diffusion of macromolecules in the brain: implications for drug delivery. Molecular pharmaceutics, 2013. 10(5): p. 1492–1504.

63. Armstrong, J.K., et al., The hydrodynamic radii of macromolecules and their effect on red blood cell aggregation. Biophysical journal, 2004. 87(6): p. 4259–4270.

64. Smith, A.J., et al., Test of the ‘glymphatic’ hypothesis demonstrates diffusive and aquaporin-4-independent solute transport in rodent brain parenchyma. eLife, 2017. 6: p. e27679.

65. Aggarwal, P., et al., Nanoparticle interaction with plasma proteins as it relates to particle biodistribution, biocompatibility and therapeutic efficacy. Adv Drug Deliv Rev, 2009. 61(6): p. 428–37.

66. Sofroniew, M.V., et al., Atrophy but not death of adult septal cholinergic neurons after ablation of target capacity to produce mRNAs for NGF, BDNF, and NT3. J Neurosci, 1993. 13(12): p. 5263–76.

67. Sofroniew, M.V., et al., Survival of adult basal forebrain cholinergic neurons after loss of target neurons. Science, 1990. 247(4940): p. 338–42.

68. Eles, J.R., et al., Meningeal inflammatory response and fibrous tissue remodeling around intracortical implants: An in vivo two-photon imaging study. Biomaterials, 2019. 195: p. 111–123.

69. Spencer, K.C., et al., Characterization of Mechanically Matched Hydrogel Coatings to Improve the Biocompatibility of Neural Implants. Scientific Reports, 2017. 7(1): p. 1952.

70. Liu, Y., et al., Soft and elastic hydrogel-based microelectronics for localized low-voltage neuromodulation. Nature Biomedical Engineering, 2019. 3(1): p. 58–68.

