## Supplementary Information for "Foreign body responses in central nervous system mimic natural wound responses and alter biomaterial functions"

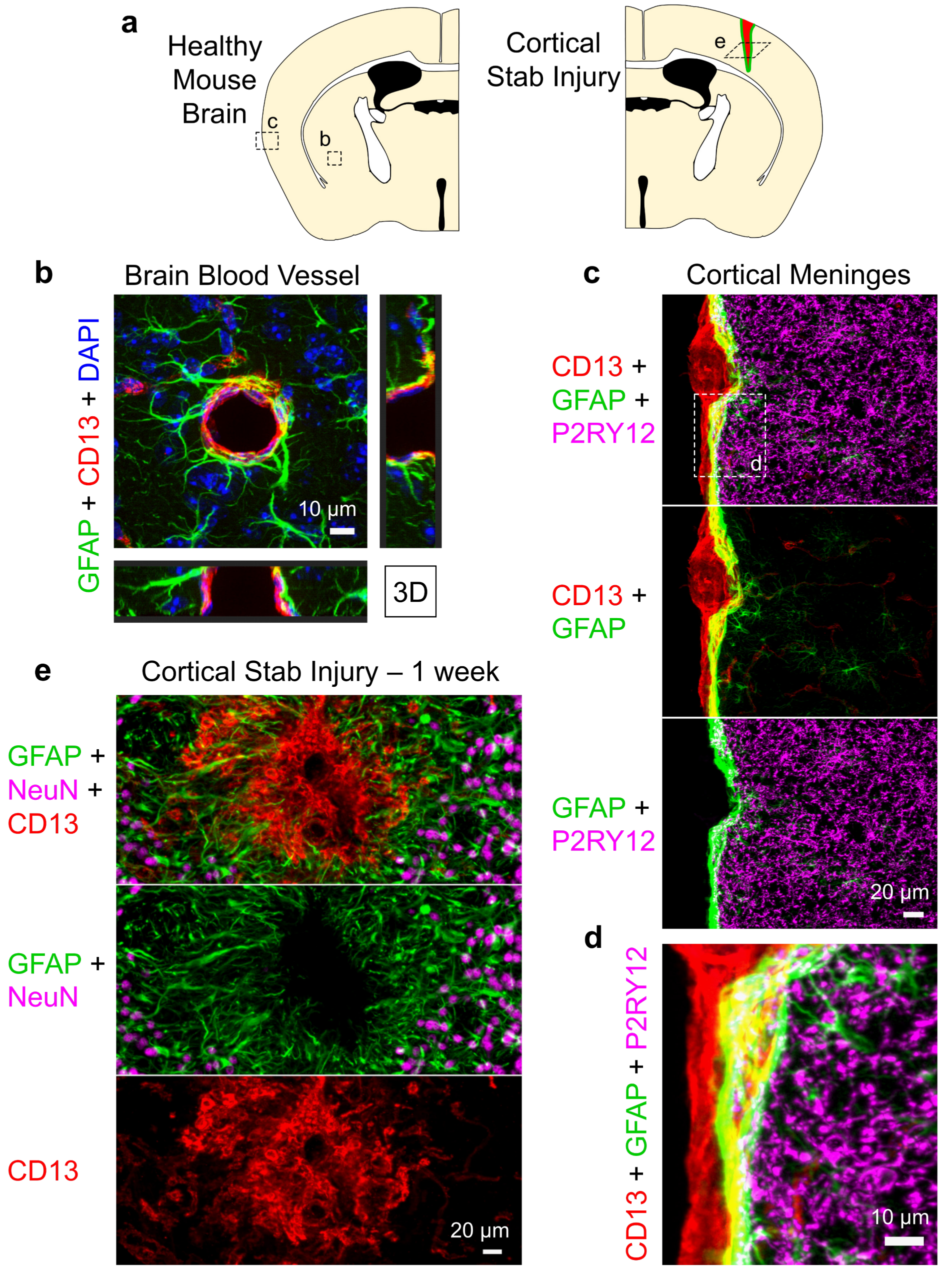


**Supplementary Figure 1 | Non-neural cells express CD13 in both the healthy and injured mouse brain and interact with border-forming astrocytes. a.** Schematics of healthy and stab injured mouse forebrain indicating locations of immunohistochemistry (IHC) images in b-e. **b.** Detail image of cerebrovasculature in mouse caudate putamen. CD13-positive pericytes along blood vessels are surrounded by cellular processes of GFAP-positive astrocytes which form glial limitans borders that segregate neural from non-neural tissue. **c. & d.** Survey and detail images of the outer surface of mouse cortex. CD13-positive stromal cells of the cerebral meninges are continually abutted by cellular processes of GFAP-positive astrocytes which form glial limitans borders that segregate neural from non-neural tissue around the entire CNS. P2RY12 staining shows microglia in healthy neural tissue. Overlapping and intertwined processes of stromal cells (red) and astrocytes (green) is seen as yellow. **e.** Detail image of a stab injury lesion in the cerebral cortex at one week after injury. CD13-positive cells accumulate at the lesion epicenter and are surrounded by the cellular processes of GFAP-positive astrocytes which segregate the CD13 non-neural compartment away from the immediately adjacent viable neural tissue compartment that contains NeuN-positive neurons.

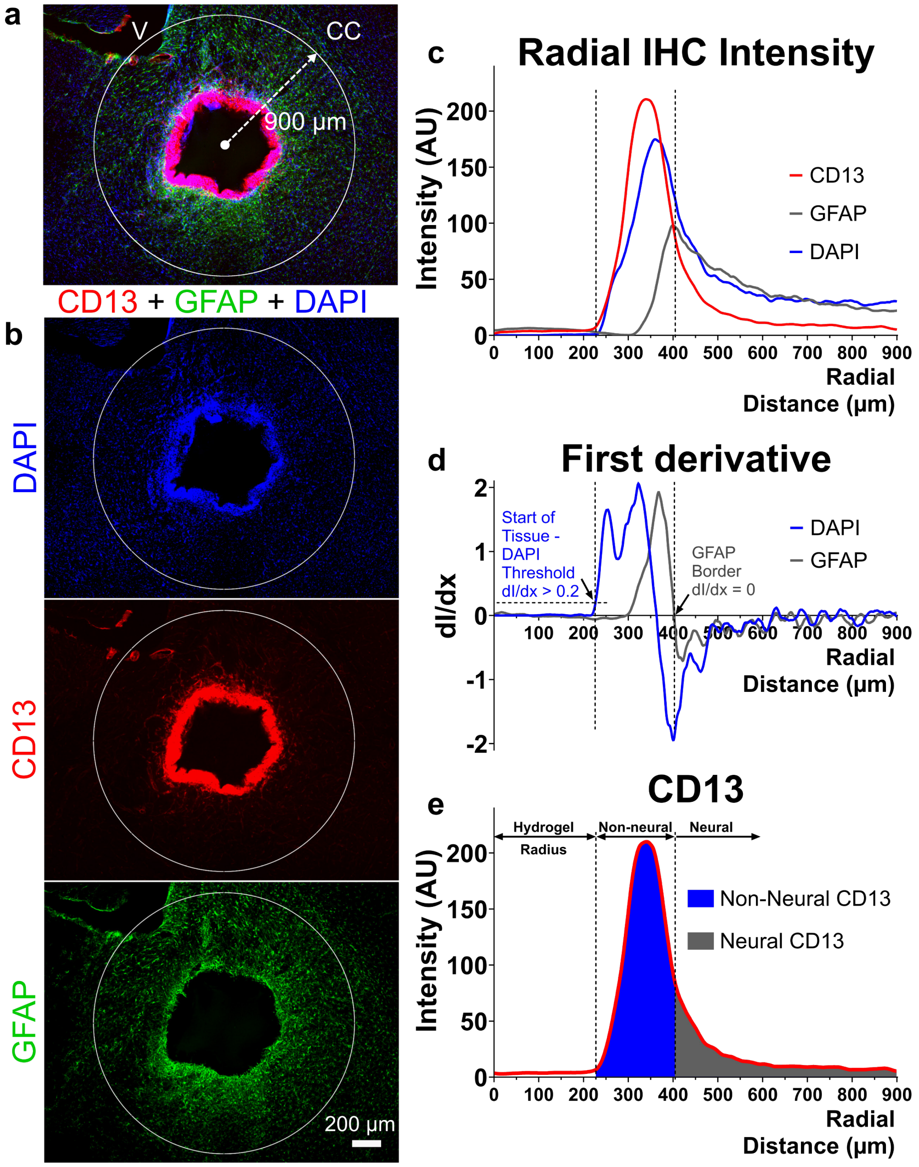


**Supplementary Figure 2 | Quantification procedure used to characterize the CNS FBR to different hydrogel formulations. a. & b.** Survey images of CD13-, GFAP- and DAPI-positive cells involved in the FBR to an example hydrogel. Hydrogel deposits were located within the caudate putamen (CP) with the ventricles (V) and corpus collosum (CC) on either side. A circular zone of analysis defined by an outer limit radius of 900 µm from the center of the hydrogel deposit was used for all samples. The zone of analysis encompassed the entire region of CP tissue around the hydrogels. **c.** Plots of average fluorescent signal intensity for the immunohistochemical (IHC) stains as a function of radial distance from the center of the hydrogel deposit. The average intensity for a specific IHC stain at a given radial distance, x, from the hydrogel deposit center was determined by averaging the intensity across the entire circumference of a circle with radius x. The average intensity was computed across the full range from 0 to 900 µm with sequential radius values increasing by a value of one pixel , where one pixel equated to 1.063 µm. **d.** Numerical differentiation methods were used to caculate the first derivative of the average IHC intensity plot for each IHC stain. The radial location of the host tissue/hydrogel interface, and hence the hydrogel radius, was defined as the radius value at which the first derivative of the DAPI intensity reached a fixed threshold of 0.2. This threshold value represented approximately 10% of the max first derivative value and was deemed to be a suitable value for detecting the start of DAPI-positive cells across a random assortment of hydrogels. The radial location of the astrocyte border was determined as the radial value where the first derviative of the GFAP intensity was zero, equating to the maxium value on the GFAP intensity plot. **e.** Area under the curve (AUC) (or definite numerical integral across a fixed range) calculations for the IHC average intensity plots were used to determine total values for either CD13 or GFAP. For CD13, the AUC value across the range between the hydrogel/tissue interface to the astrocyte border was defined as the total non-neural CD13 value while the CD13 intensity from the astrocyte border to the edge of the analysis region was defined as the neural tissue CD13 value. For total GFAP, a range from the hydrogel-tissue interface to the edge of the analysis region was assessed.

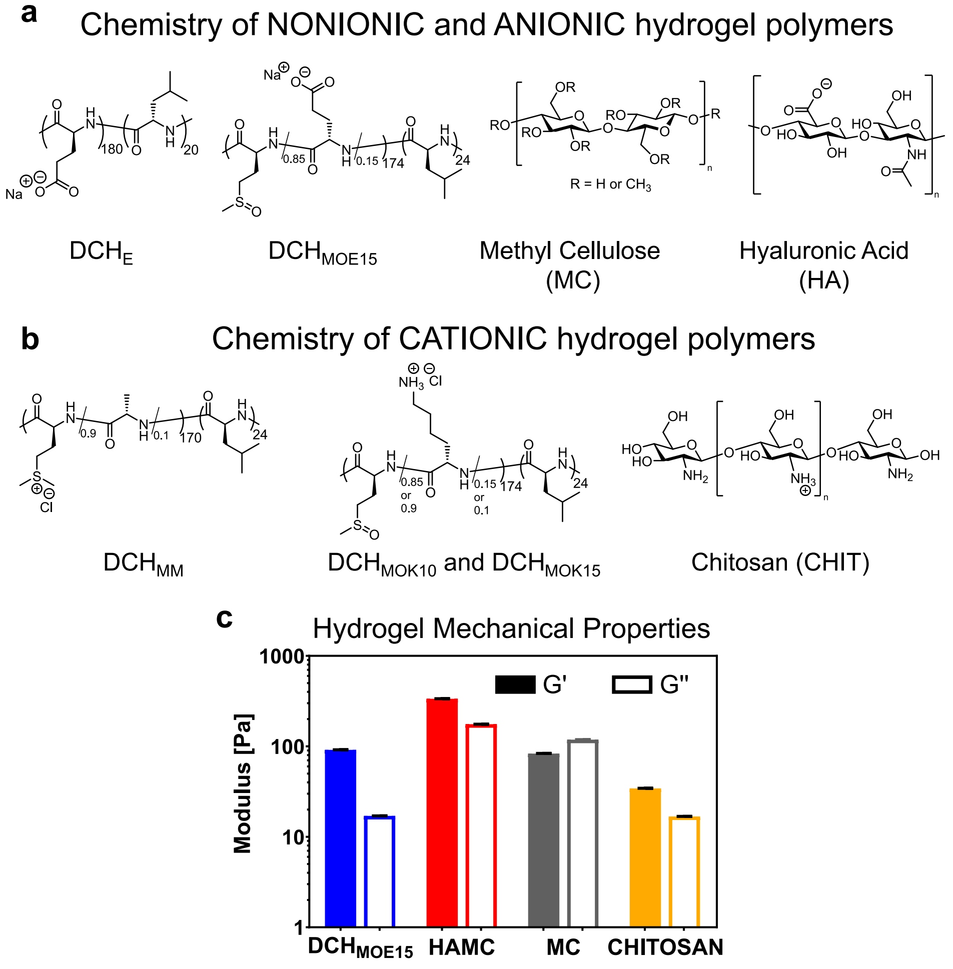


**Supplementary Figure 3 | Chemical structures of different polymers used to formulate hydrogels evaluated in this study. a.** Non-ionic and anionic polymers **b.** Cationic polymers. Note that $DCH_{MOE15}$, $DCH_{MOK10}$ and $DCH_{MOK15}$ consisted of $DCH_{MO}$ to which small amounts of either negatively charged glutamate (E), or positively charged lysine (K) had been incorporated into the hydrophilic polypeptide block as a statistical copolymer. We also modified the poly(L-methionine-*stat*-L-alanine) segments in the parent polymer used to formulate $DCH_{MO}$ by selective methylation of methionine residues to form the cationic methyl-sulfonium based $DCH_{MM}$ [9]. **c.** Mechanical properties of some of the hydrogels used in this study as evaluated by dynamic rheology. Mechanical properties of the other DCH used are detailed in our previous publications [3, 9, 26].

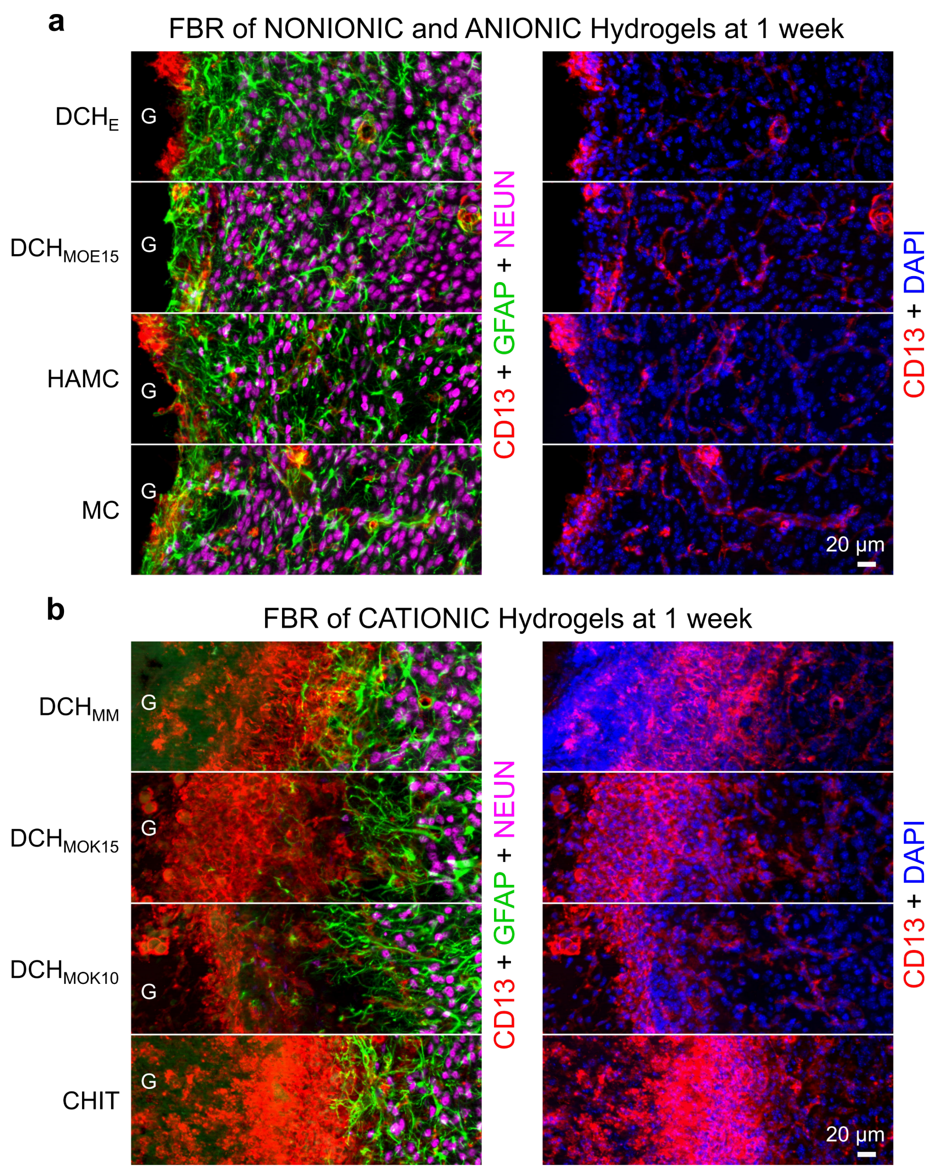


**Supplementary Figure 4 | Detail images of different hydrogel FBRs at one week after injection into mouse caudate putamen (CP). a.** Nonionic and anionic hydrogels exhibit barely detectable levels of CD13-positive cells at host interfaces. **b.** Cationic hydrogels exhibit large rims of CD13-positive cells infiltrating into deposits at their interfaces with host tissue.

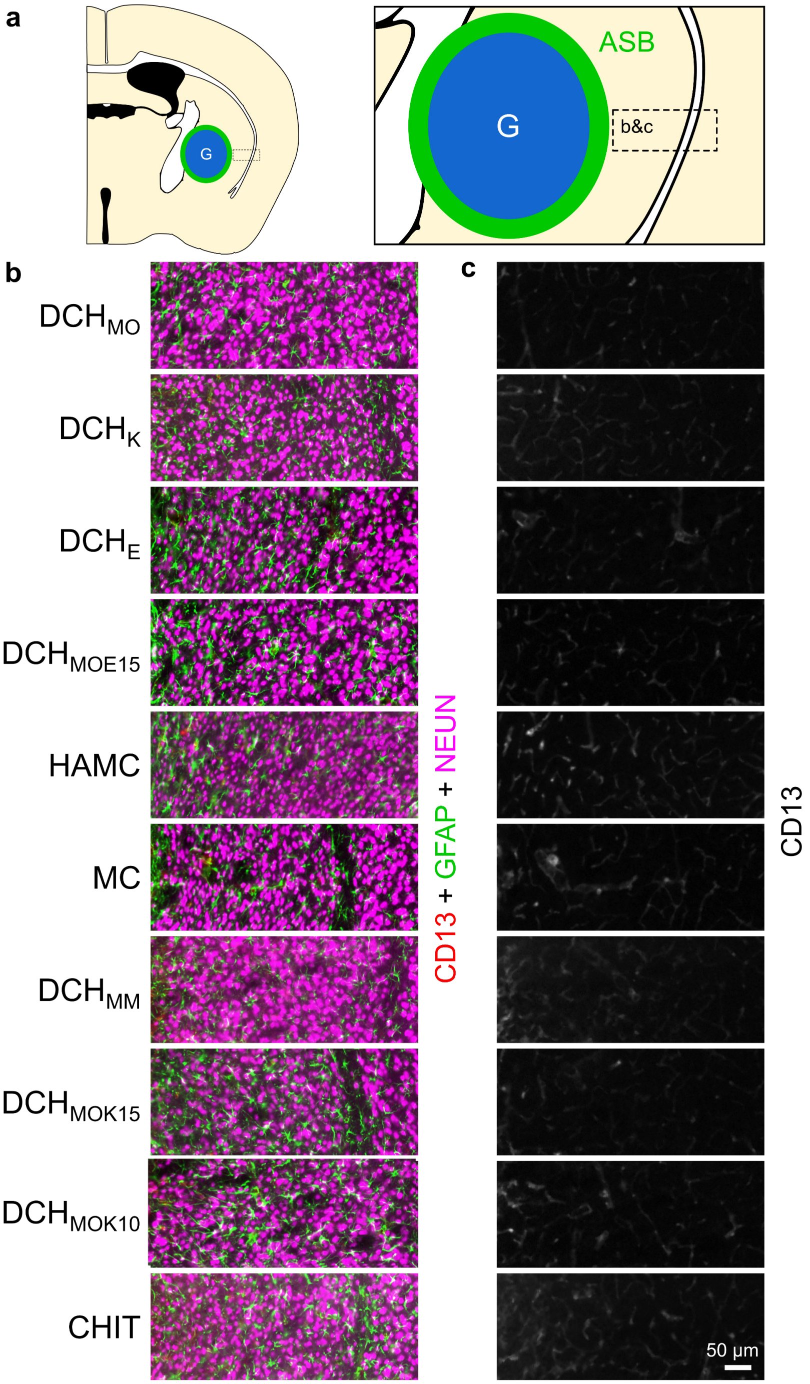


**Supplementary Figure 5 | Detail images of neural tissue immediately adjacent to the astrocyte border (ASB) along hydrogel interfaces for the different hydrogels tested. a.** Schematic of mouse forebrain indicating the location of the images in b. & c. **b.** The appearance and density of GFAP-positive astrocytes and NeuN-positive neurons in the preserved neural tissue adjacent to hydrogels was not detectably different across all samples regardless of the differences in severity of FBR in the nearby non-neural tissue. **c.** CD13-positive cells in the preserved neural tissue were similar in appearance to those in uninjured tissue and were limited to normally appearing perivascular cells across all hydrogels tested regardless of the severity of the FBR in the nearby non-neural tissue.

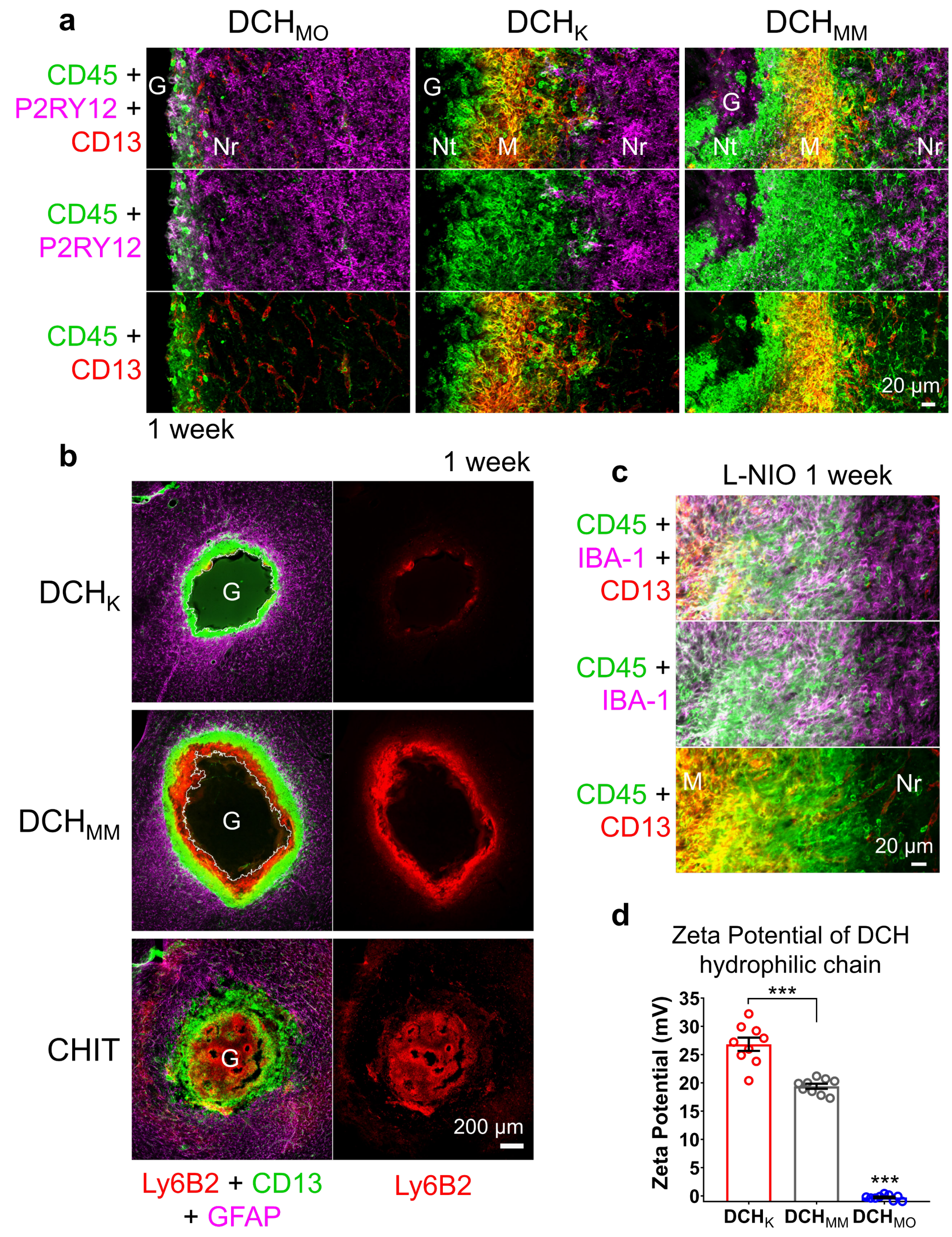


**Supplementary Figure 6 | Characterization of inflammatory cells involved in hydrogel FBR or CNS wound response. a.** Detail images of the escalating recruitment of peripherally derived inflammatory cells by $DCH_{MO}$, $DCH_{K}$, $DCH_{MM}$ at one week after injection. For all hydrogels (G), the neural tissue (Nr) immediately adjacent to deposits contains reactive, CNS-derived microglia that are Iba-1-positive, P2RY12-positive and CD13-negative. Between neural tissue and hydrogels there are zones of infiltrating macrophages (M) that are CD13-positive and CD45-positive and P2RY12-negative and that increase in thickness and intensity from $DCH_{MO}$ to $DCH_{K}$ and $DCH_{MM}$. Between the layer of macrophages and $DCH_{K}$ and $DCH_{MM}$ there are increasing levels of CD45-positive, CD13-negative and P2YR12-negative cells that stain for Ly6B2 (see panel b) and are neutrophils (Nt). **b.** Survey images of different cationic hydrogels comparing infiltration Ly6B2-postive neutrophils at one week after injection. Note for $DCH_{K}$, the sparse number of neutrophils (red) between the layer CD-13 positive macrophages (green) and gel deposit, in contrast with the thick layer of neutrophils in $DCH_{MM}$ and the robust neutrophil invasion of the entire center of the chitosan (CHIT) deposit. **c.** Detail images of an L-NIO induced stroke lesion at one week after injection. The recruitment of inflammatory cells to a stroke lesion occurs in a similar manner to that seen for hydrogel with formation of a non-neural lesion core of predominantly stromal cells and macrophages surrounded by surviving neural tissue. **d.** Zeta Potential measurements for the hydrophilic chain of DCH showing that $DCH_{K}$ has a greater overall cationic charge compared to $DCH_{MM}$ because the latter is a statistical copolymer that includes 10% uncharged alanine. $DCH_{MO}$ has essentially zero charge. ***$P < 0.0001$ versus $DCH_{K}$, one-way ANOVA with Bonferroni. Graph shows mean ± s.e.m with individual data points showing *n* replicate samples per group.

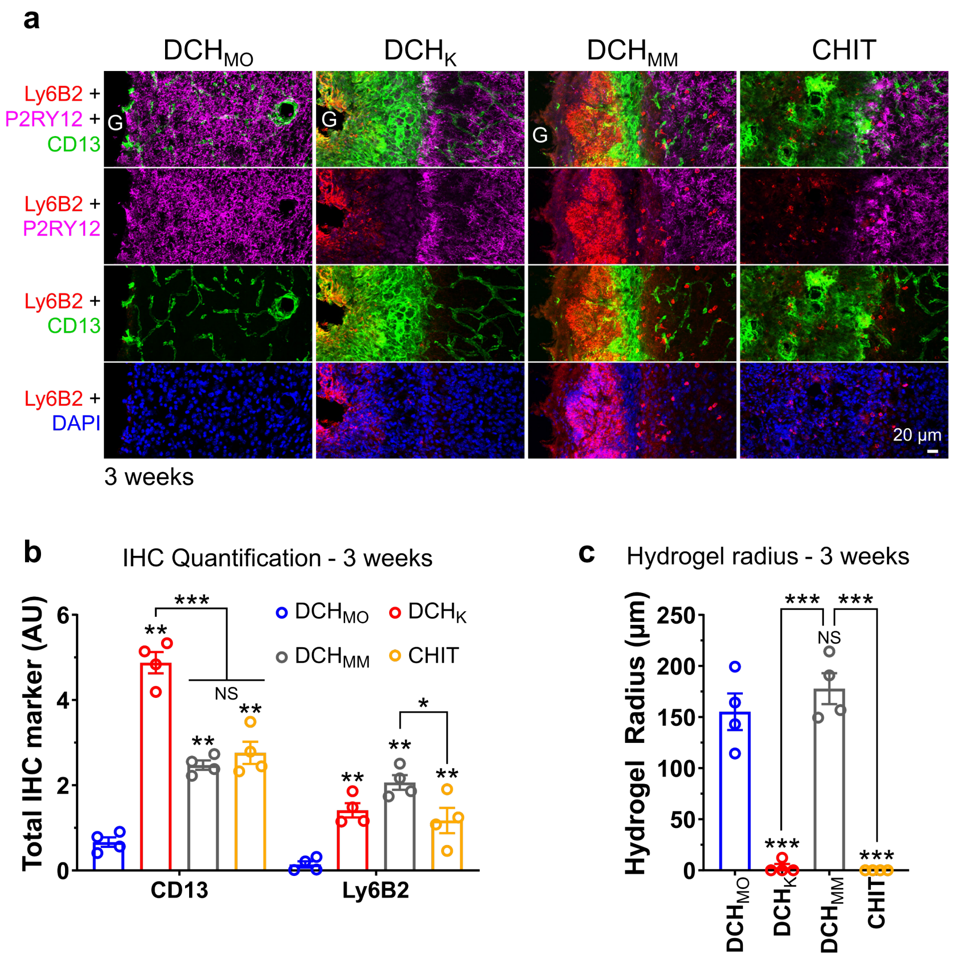


**Supplementary Figure 7 | Characterization of inflammatory cells involved in hydrogel FBR at 3 weeks after injection. a.** Detail images of hydrogel-tissue interface showing the distribution of neutrophils (Ly6B2), macrophages (CD13) and microglia (P2RY12) at 3 weeks for different hydrogels. Persistence of Ly6B2-positive neutrophils at the cationic $DCH_{MM}$ hydrogel surface at three weeks after injection is apparent. $DCH_K$ and chitosan (CHIT) hydrogels show infiltration of macrophages into hydrogels at 3 weeks. **b.** Quantification of total CD13 and Ly6B2 staining for hydrogels at three weeks after injection. **P < 0.007 for all hydrogels versus $DCH_{MO}$ for both CD13 and Ly6B2, ***P < 0.0001 for $DCH_{MM}$ and CHIT versus $DCH_K$ for CD13, Not Significant (NS) and *P < 0.02 for $DCH_{MM}$ versus CHIT for CD13 and Ly6B2, two-way ANOVA with Bonferroni. **c.** Quantification of hydrogel radius for the different materials at 3 weeks. $DCH_K$ and CHIT are mostly resorbed at the 3-week time point. $DCH_{MM}$ is not consumed despite prevalent inflammatory cells at the hydrogel interface indicating frustrated phagocytosis. $DCH_{MO}$ is also not resorbed but shows limited peripheral inflammatory cells at the hydrogel interface. NS not significant and ***P < 0.0001 for all hydrogels versus $DCH_{MO}$ and ***P < 0.0001 for $DCH_K$ and CHIT versus $DCH_{MM}$, one-way ANOVA with Bonferroni. Graphs show mean ± s.e.m with individual data points showing *n* mice per group.

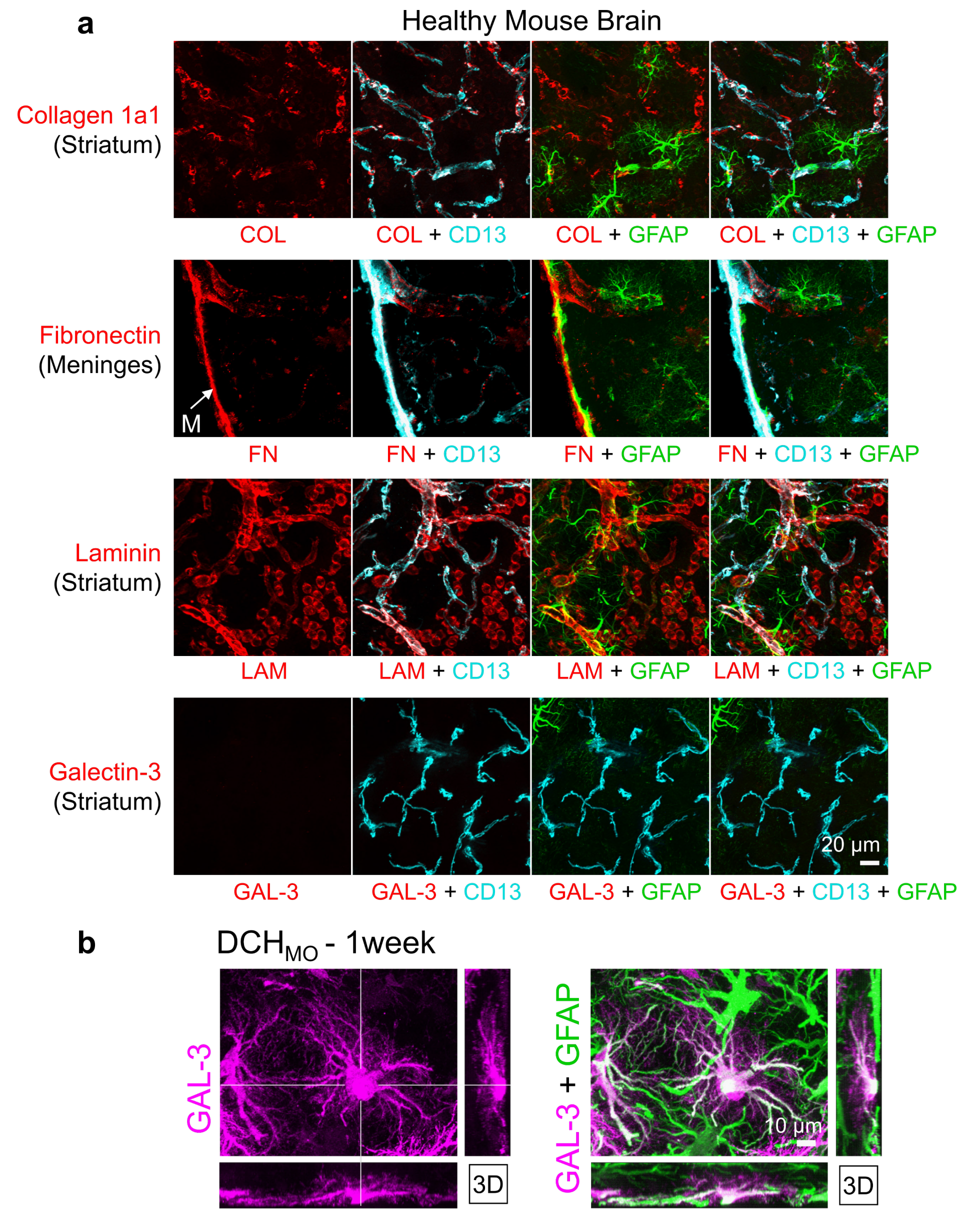


**Supplementary Figure 8 |Characterization of fibrosis and stromal cell elements in healthy and hydrogel injected CNS. a.** Detail images of normal wildtype mouse brain stained for extracellular matrix and stromal cell associated markers collagen 1a1, fibronectin, laminin and galectin-3. CD13-positive cells along blood vessels are also positive for collagen-1a1, fibronectin and laminin positive. CD13-positive cells along meninges (M) are also positive for fibronectin. Galectin-3 is undetectable in normal tissue. **b.** High magnification 3D view shows that galectin-3 is robustly expressed by some but not all reactive astrocytes near the $DCH_{MO}$ interface at one week. GFAP and galectin-3 colocalization is seen as white staining. Galectin-negative astrocytes are stained only green.

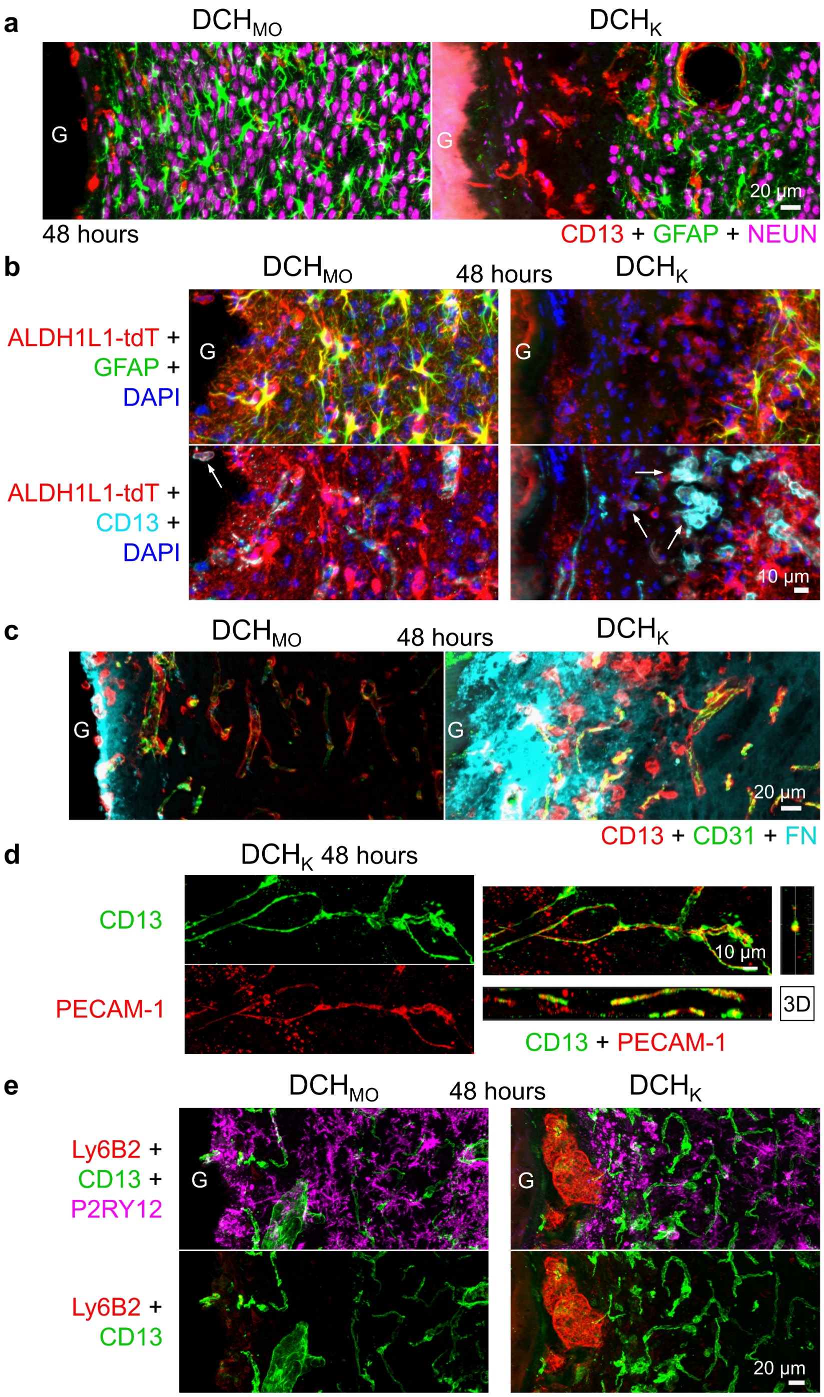


**Supplementary Figure 9 | Characterization of hydrogel evoked tissue damage. a.** Comparison of the tissue interface for $DCH_{MO}$ and $DCH_K$ at 48 hours. Note the clear zone of neural tissue loss in $DCH_K$ indicated by loss of GFAP and NeuN positive cells and presence of CD13 cells, whereas such tissue loss is mostly absent for $DCH_{MO}$. **b.** Hydrogel induced loss of astrocytes in Aldh1l1-tdT mice results in tdT reporter debris within the damaged tissue zone that is phagocytosed by CD13-positive macrophages (white arrows). **c.** Neural tissue damage at the hydrogel interface results in extracellular deposition of extravasated blood-borne fibronectin, which is more apparent in $DCH_K$ compared to $DCH_{MO}$. **d.** CD13 and PECAM-1 positive thin vessels in the zone of tissue damage at the $DCH_K$ tissue interface. **e.** Ly6B2 positive neutrophils infiltrate the region of tissue damage that occurs at the interface of host tissue with $DCH_K$, but not at the interface with $DCH_{MO}$.

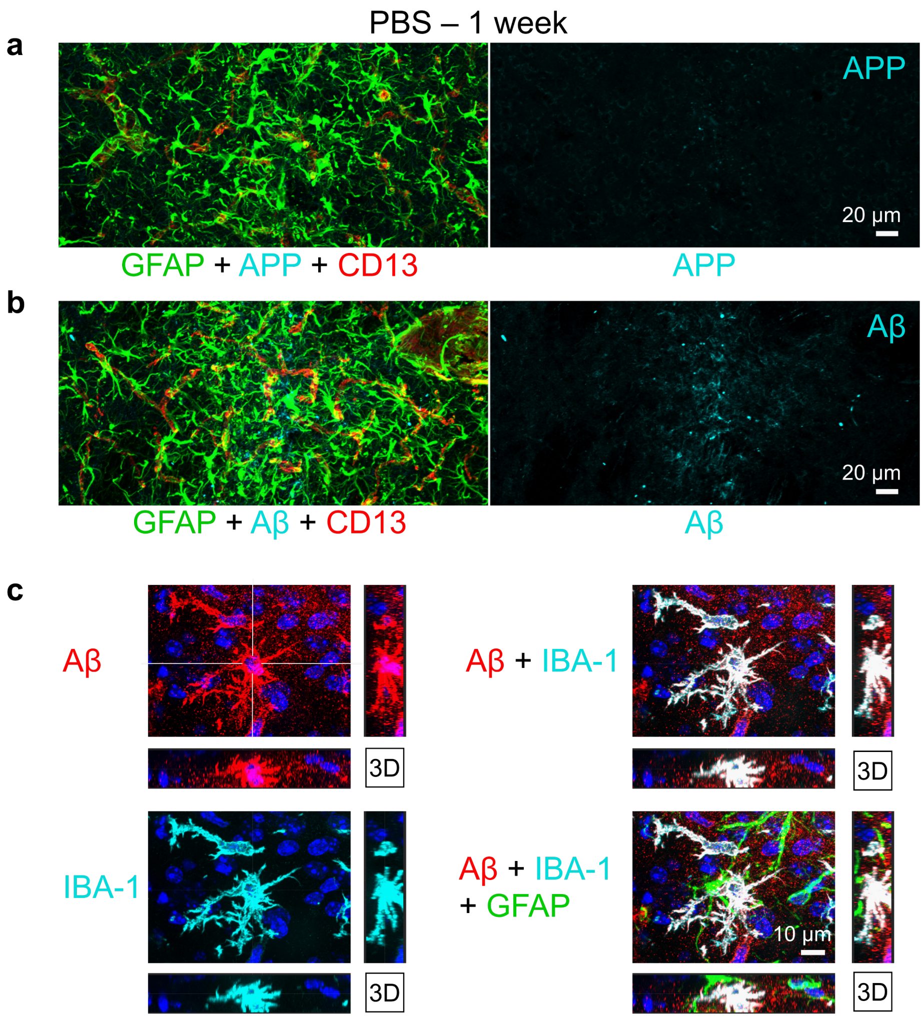


**Supplementary Figure 10 | Characterization of APP and Aβ after injection of PBS or hydrogel. a,b.** Infusions of PBS cause limited axonal damage, APP accumulation (**a**) and Aβ formation (**b**) only in narrow zone of damage caused by the needle track. **c.** After injection of $DCH_K$, Aβ is colocalized prominently within IBA-1-positive microglia but does not colocalize with GFAP-positive astrocytes.

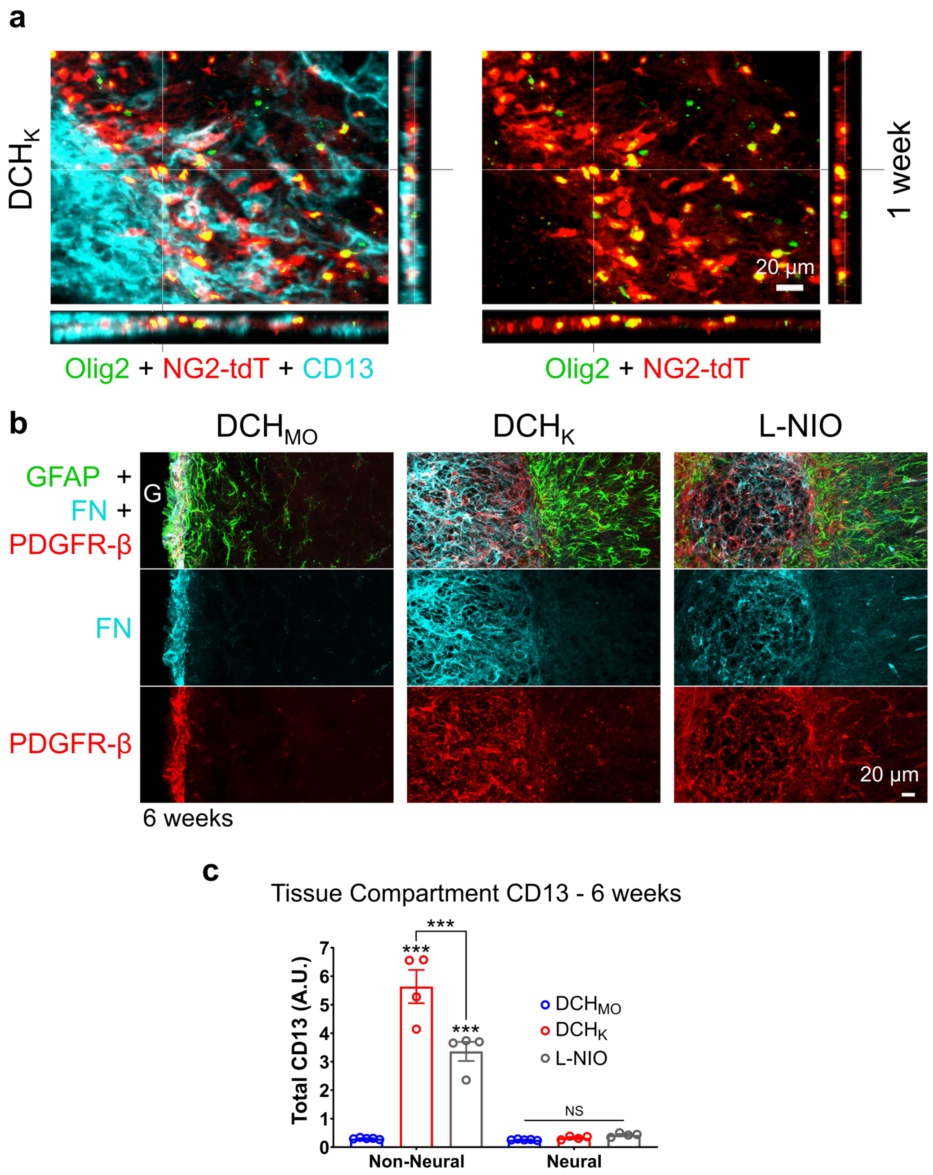


**Supplementary Figure 11 | Characterization of glia and non-neural cells in chronic FBR to hydrogels. a.** Detail images showing that OPCs positive for Olig2 and NG2-targeted reporter protein (NG2-tdT) remain partitioned in the neural tissue compartment during the FBR to hydrogels. **b.** Detail images showing fibronectin (FN) and PDGFR-β in the chronic fibrotic tissue abutting the mature GFAP-positive astrocyte limitans border at 6 weeks after $DCH_{MO}$, $DCH_K$ and L-NIO stroke. **c.** Quantification of total CD13 staining in non-neural and neural tissue compartments for $DCH_{MO}$, $DCH_K$ and L-NIO stroke after 6 weeks ***$P < 0.0001$ versus $DCH_{MO}$ and NS for neural compartment for all samples, two-way ANOVA with Bonferroni. Graph shows mean ± s.e.m with individual data points showing *n* mice per group.

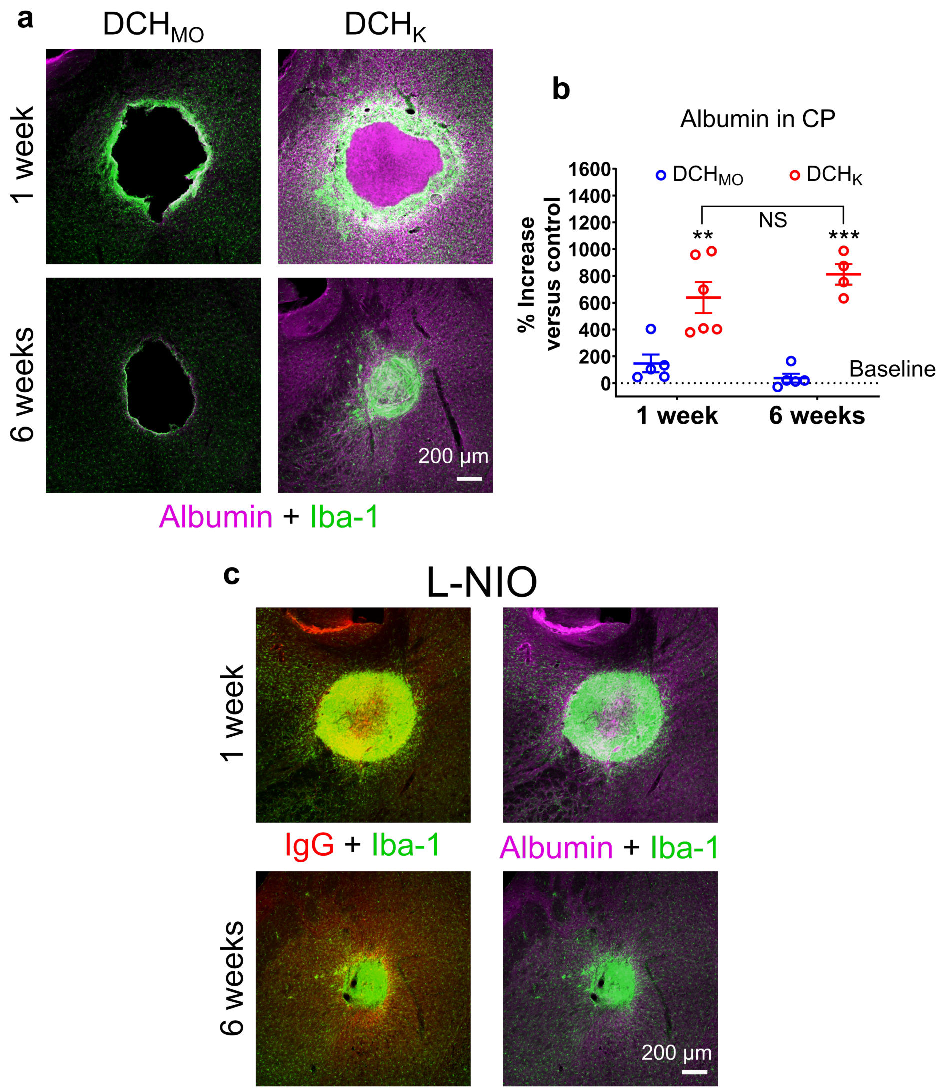


**Supplementary Figure 12 | Analysis of blood-brain barrier (BBB) leak and repair following hydrogel infusions or L-NIO-induced stroke and formation of the astrocyte limitans border. a.** Survey images of albumin penetration into neural parenchyma at one week and six weeks after DCH$_{MO}$ and DCH$_K$ injections. **b.** Quantification of the percentage increase in albumin levels in the hydrogel injected caudate putamen (CP) normalized to the non-injected contralateral side. **P < 0.001 and ***P < 0.0001 for DCH$_K$ versus DCH$_{MO}$ at 1 week and 6 weeks respectively and NS for DCH$_K$ samples between the two time points, two-way ANOVA with Bonferroni. Graph shows mean ± s.e.m with individual data points showing *n* mice per group. **c.** Survey images of penetration serum proteins albumin and IgG into neural parenchyma at one and six weeks after L-NIO-induced strokes.

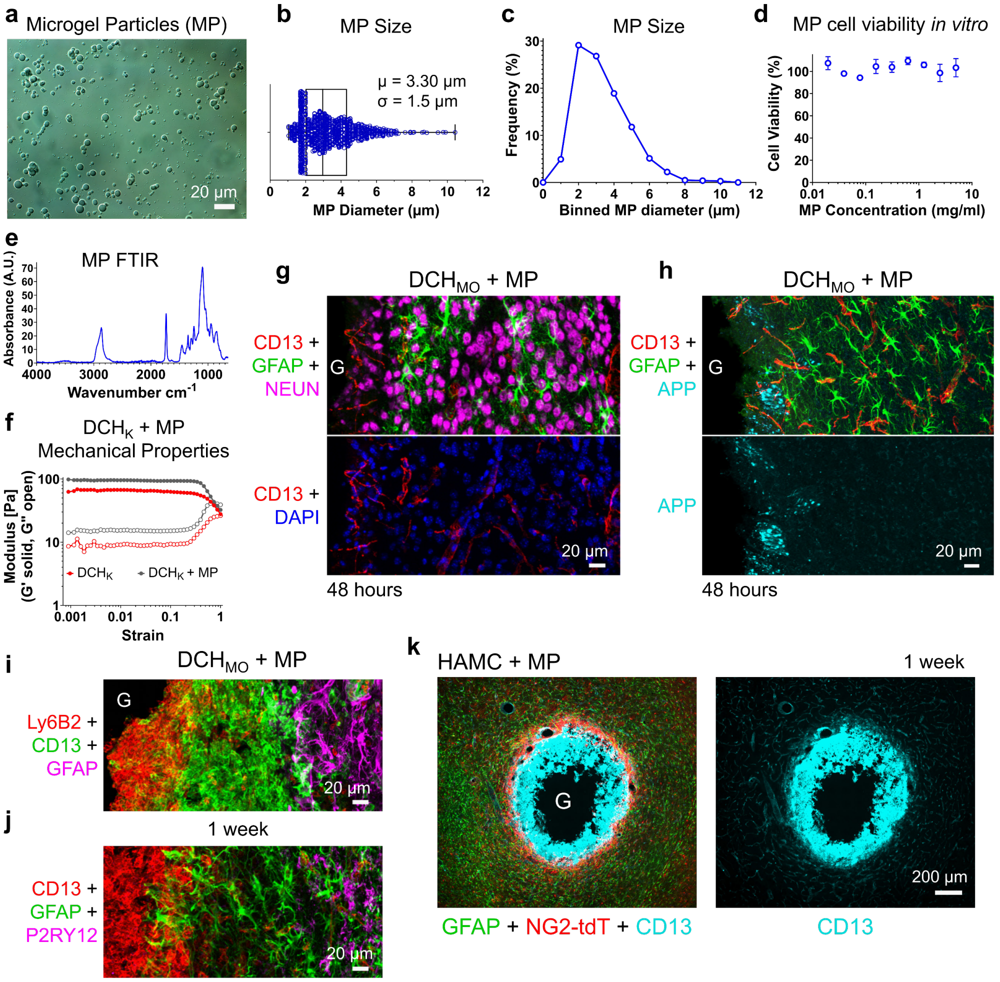


**Supplementary Figure 13 | Characterization of PEG microgel particles (MP) and their effect on the FBR when combined with physical hydrogels. a.** Differential interference contrast (DIC) microscopy image of PEG microgel particles (MP) formed by inverse emulsion. **b,c.** Quantification of MP diameter spread and frequency distribution. **d.** Cell viability of cultured neural progenitor cells exposed to MP for twenty-four hours at various concentrations up to 5 mg/ml. MP are non-toxic to neural progenitors in culture at all concentrations tested **e.** ATF-FTIR spectrum of dried MP. The ester stretch (≈1730 $cm^{-1}$) and the loss of the acrylate C=C stretch (1675–1600 $cm^{-1}$) were used to confirm presence of the covalent crosslinked network. **f.** Characterization of mechanical properties of DCH loaded with MP shows that MPs impart a slight stiffening but no effect on the injectability of the hydrogel at high strain. **g,h.** Detail images showing that inclusion of MP into $DCH_{MO}$ has no detectable adverse effects on neural tissue (g) and does not detectably increase axonal injury as identified by APP accumulation (h). **i.** Detail image showing a pronounced infiltration of CD13-positive macrophages and Ly6B2-positive neutrophils at 1 week within deposits of $DCH_{MO}$ hydrogels loaded with MP. **j.** Detail image showing that P2Y12R-positive microglia do not infiltrate into $DCH_{MO}$ + MP deposits. **k.** Detail image showing that loading MP into HAMC hydrogels also leads to a pronounced recruitment and infiltration of CD13-positve cells by 1 week and that GFAP-positive astrocytes and NG2-tdT-positive OPCs do not migrate into CD13 positive cell zones.

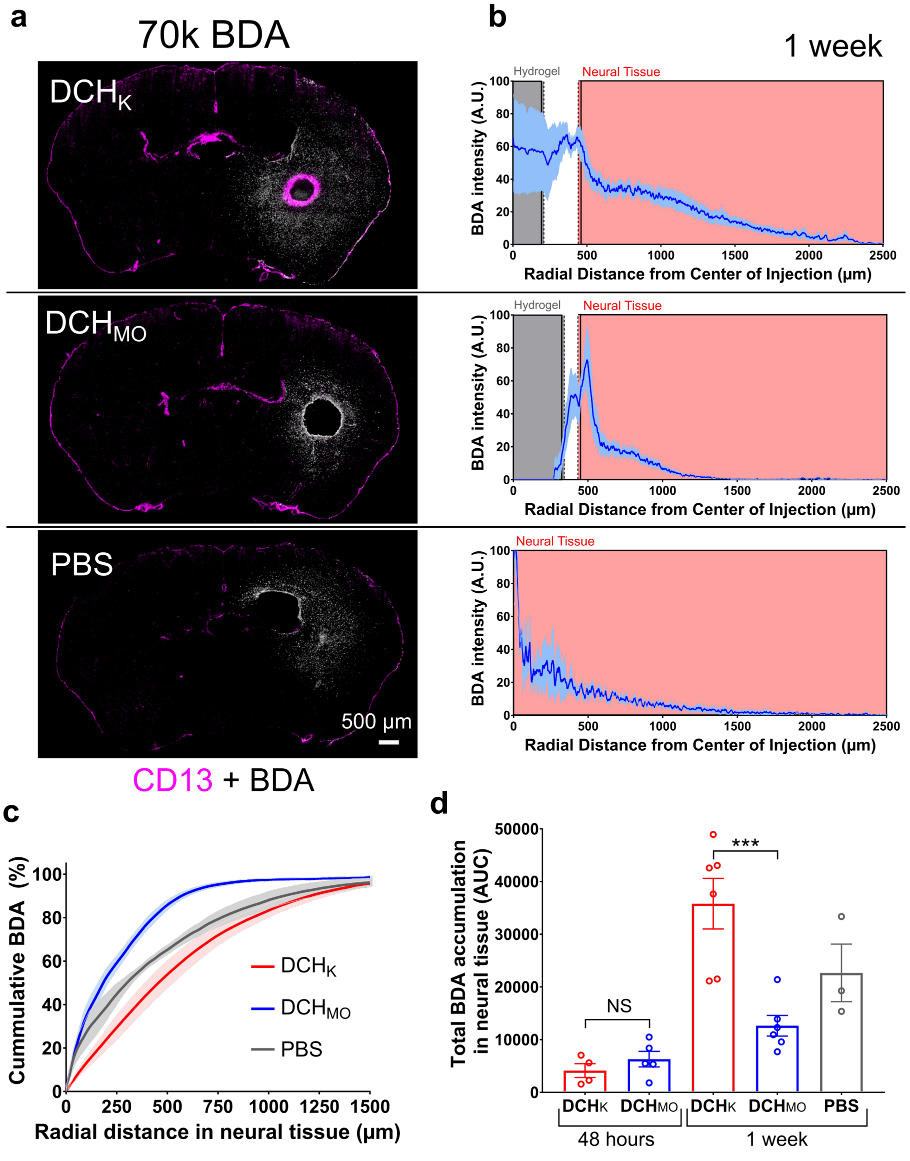


**Supplementary Figure 14 | Analysis of biodistribution of 70kDA BDA (BDA-70) released from hydrogels in the striatum. a.** Survey images of BDA-70 delivery using $DCH_K$, $DCH_{MO}$, and PBS carriers at 1 week. **b.** BDA intensity plots of BDA-70 released into the lateral striatum and cortex as measured radially from the center of the hydrogel injection. Grey shaded regions indicate location of the hydrogel, red shaded regions indicate neural parenchyma tissue, the white unshaded zone between the grey and red shaded regions includes non-neural tissue and start of the astrocyte limitans border. Data are mean ± s.e.m, with s.e.m. represented as light blue shaded areas around the average intensity plot for each condition. **c.** Cumulative BDA-70 accumulation in neural tissue at 1 week for the hydrogel and PBS carriers. **d.** Total BDA-70 accumulation for the different conditions at 48 hours and 1 week. There was no difference in burst release from the two hydrogels but $DCH_K$ showed increased accumulation of BDA-70 in neural tissue compared to $DCH_{MO}$ at 1 week. NS and ***$P < 0.0001$ for $DCH_K$ versus $DCH_{MO}$, two-way ANOVA with Bonferroni. Data are mean ± s.e.m with individual data points showing *n* mice per group.

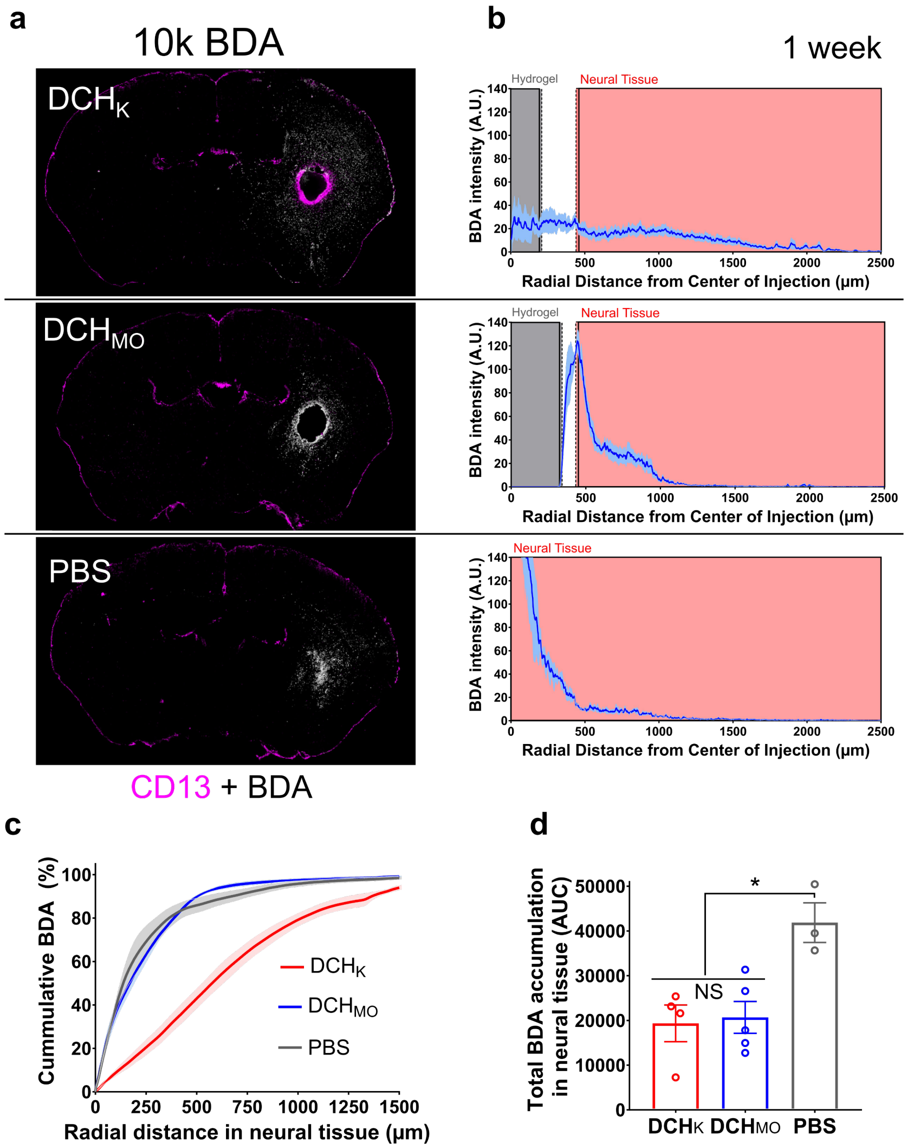


**Supplementary Figure 15 | Analysis of biodistribution of 10kDA BDA (BDA-10) released from hydrogels in the striatum. a.** Survey images of BDA-10 delivery using $DCH_K$, $DCH_{MO}$, and PBS carriers at 1 week. **b.** BDA intensity plots of BDA-10 released into the lateral striatum and cortex as measured radially from the center of the hydrogel injection. Data are mean ± s.e.m, with s.e.m. represented as light blue shaded areas around the average intensity plot for each condition**. c.** Cumulative BDA-10 accumulation in neural tissue at 1 week measured radially for the hydrogel and PBS carriers. $DCH_K$ shows significantly greater depth of BDA-10 penetration into neural parenchyma compared to $DCH_{MO}$. **d.** Total BDA-10 accumulation for the different conditions at 1 week shows no difference between the hydrogels but a small increased accumulation for the PBS carrier. *$P < 0.02$ for PBS versus $DCH_{MO}$ or $DCH_K$, one-way ANOVA with Bonferroni. Data are mean ± s.e.m with individual data points showing *n* mice per group.

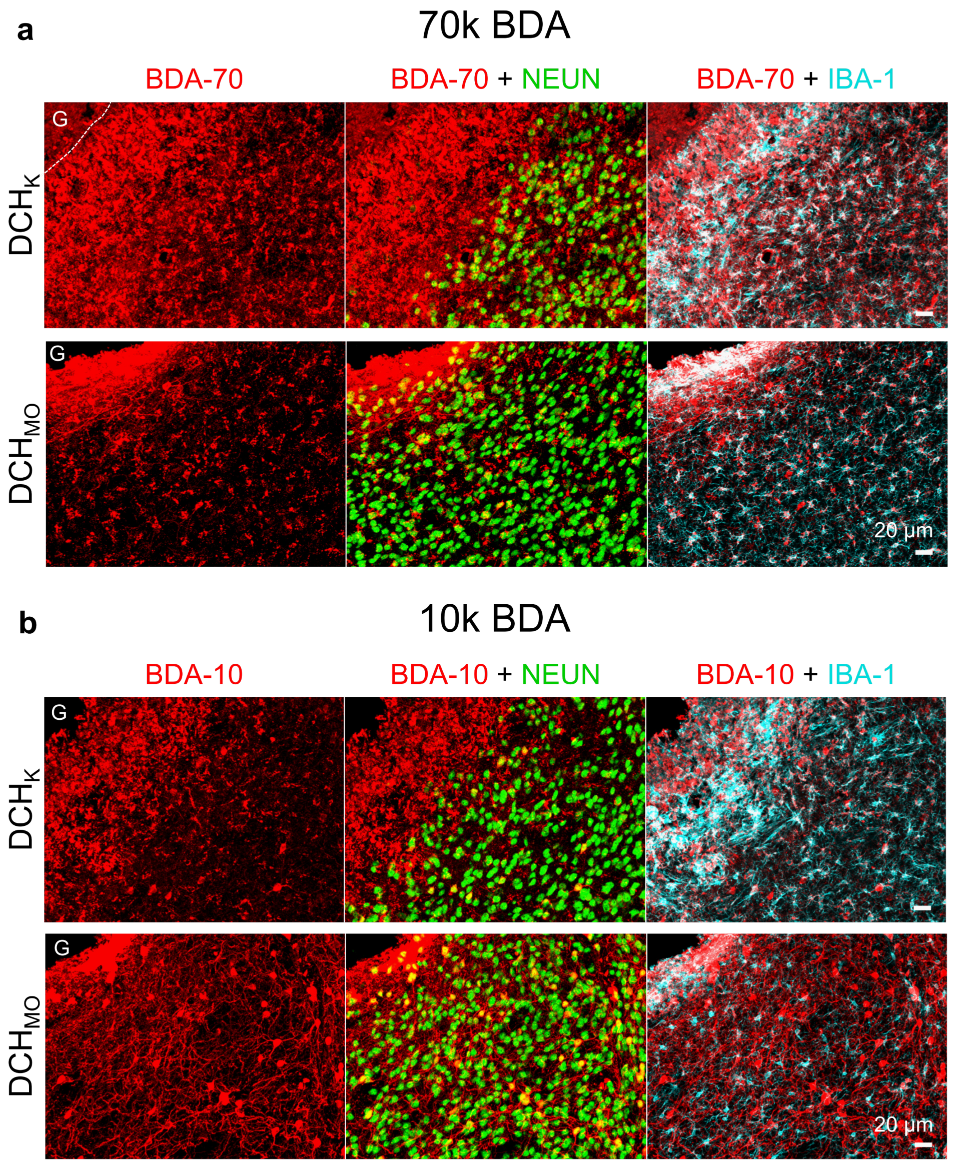


**Supplementary Figure 16 | BDA delivered from hydrogels is accumulated in neurons, microglia and macrophages in a size and hydrogel dependent manner. a.** Detail images showing the relative uptake of released BDA-70 by NeuN-positive neurons and IBA-1 positive microglia/macrophages at 1 week for DCH$_K$ and DCH$_{MO}$. **b.** Detail images showing the relative uptake of released BDA-10 by NeuN-positive neurons and IBA-1 positive microglia/macrophages at 1 week for DCH$_K$ and DCH$_{MO}$.

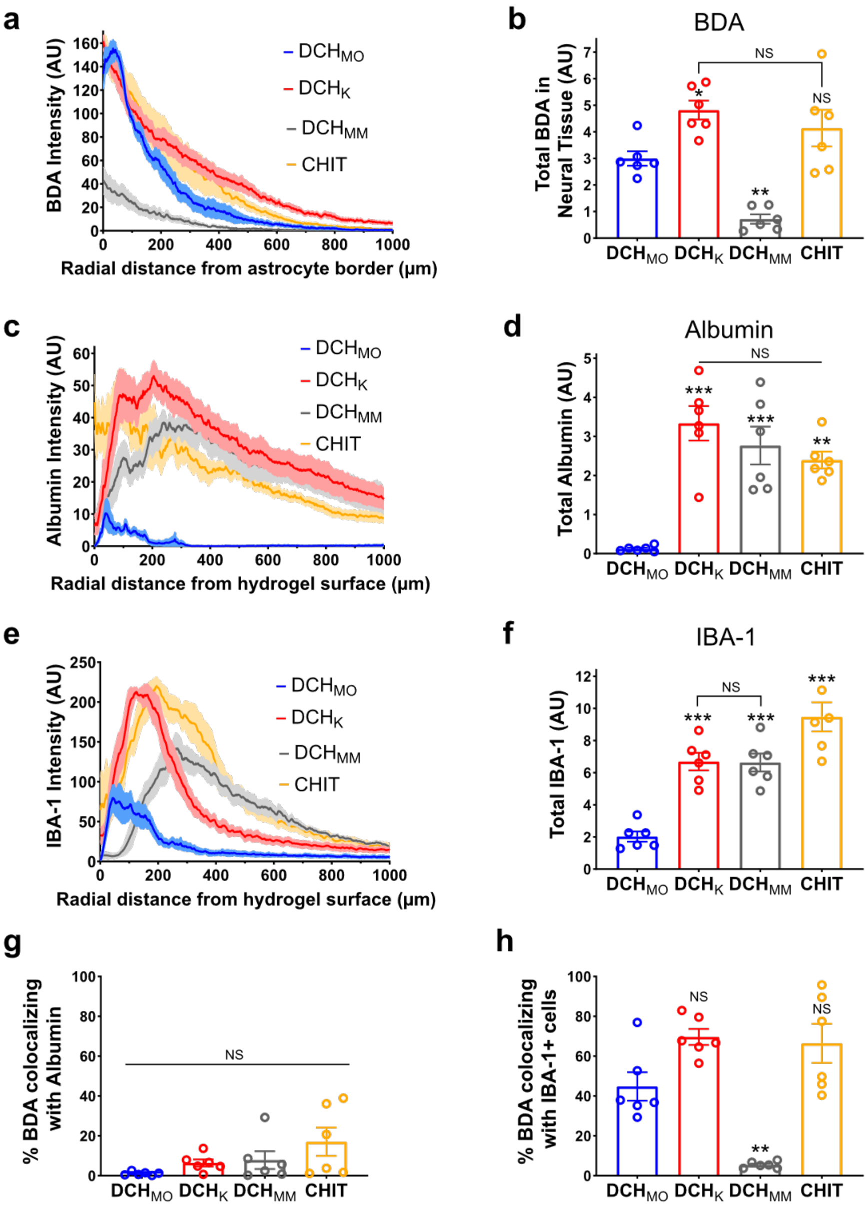


**Supplementary Figure 17 | Quantification of BDA delivery and FBR for hydrogels at 2 weeks. a.** BDA intensity distribution within neural parenchyma for the different hydrogels. **b.** Total BDA accumulation in neural tissue for the different hydrogel carriers. NS Not Significant, $^{*}P < 0.05$, $^{**}P < 0.005$ for hydrogels versus DCH$_{MO}$. **c.** Average albumin intensity profile for the different hydrogels. **d.** Total albumin staining around hydrogel deposits. $^{***}P < 0.0002$ for DCH$_{K}$ and DCH$_{MM}$ versus DCH$_{MO}$, $^{**}P<0.001$ for CHIT versus DCH$_{MO}$, NS for comparisons between DCH$_{K}$, DCH$_{MM}$, and CHIT. **e.** Average intensity plot for Iba-1 IHC staining measured radially from the hydrogel surface **f.** Total Iba-1 around hydrogel deposits. $^{***}P < 0.0002$ for DCH$_{K}$, DCH$_{MM}$ or CHIT versus DCH$_{MO}$. **g.** Percentage of total BDA colocalizing with albumin for the different hydrogels. There was no significant colocalization of BDA-10 with albumin for all samples **h.** Percentage of total BDA-10 colocalizing with Iba-1 for the different hydrogels. $^{**}P < 0.002$ for DCH$_{MM}$ versus DCH$_{MO}$ and NS for DCH$_{K}$ or CHIT versus DCH$_{MO}$. All statistical analyses are one-way ANOVA with Bonferroni. In a., c. and e. figures the data are mean ± s.e.m (lighter shaded area) for n=6 mice per group. Data in figures b., d. and f-h. are mean ± s.e.m with individual data points showing *n* mice per group.

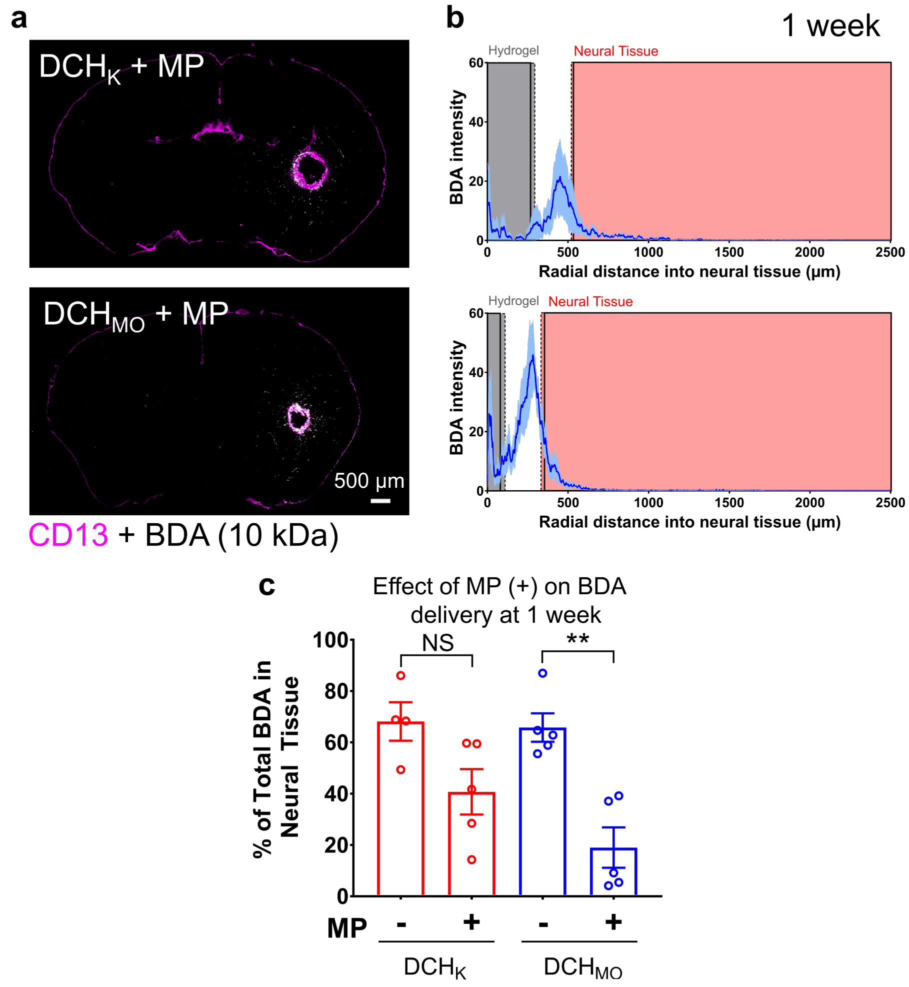


**Supplementary Figure 18 | BDA delivered from microgel particles (MP) suspended in hydrogels is affected by MP dependent FBR. a.** Survey images of BDA-10 biodistribution at 1 week delivered from MP that were suspended in DCH$_K$ and DCH$_{MO}$. **b.** Intensity plots of BDA-10 released from MP loaded hydrogels. Grey shaded regions indicate location of the hydrogel, red shaded regions indicate neural parenchyma tissue. Data are mean ± s.e.m, with s.e.m. represented as light blue shaded areas around the average intensity plot for each condition. **c.** Comparisons of the Percentage of total BDA accumulated in neural tissue at 1 week for DCH$_K$ and DCH$_{MO}$ when delivered with (+) or without (-) MP. For DCH$_{MO}$ there was a significant decrease in the amount of BDA-10 delivered to neural parenchyma with the use of the MP, ***$P < 0.003$ for DCH$_{MO}$ + MP versus DCH$_{MO}$ only, one-way ANOVA with Bonferroni. Data are mean ± s.e.m with individual data points showing *n* mice per group.

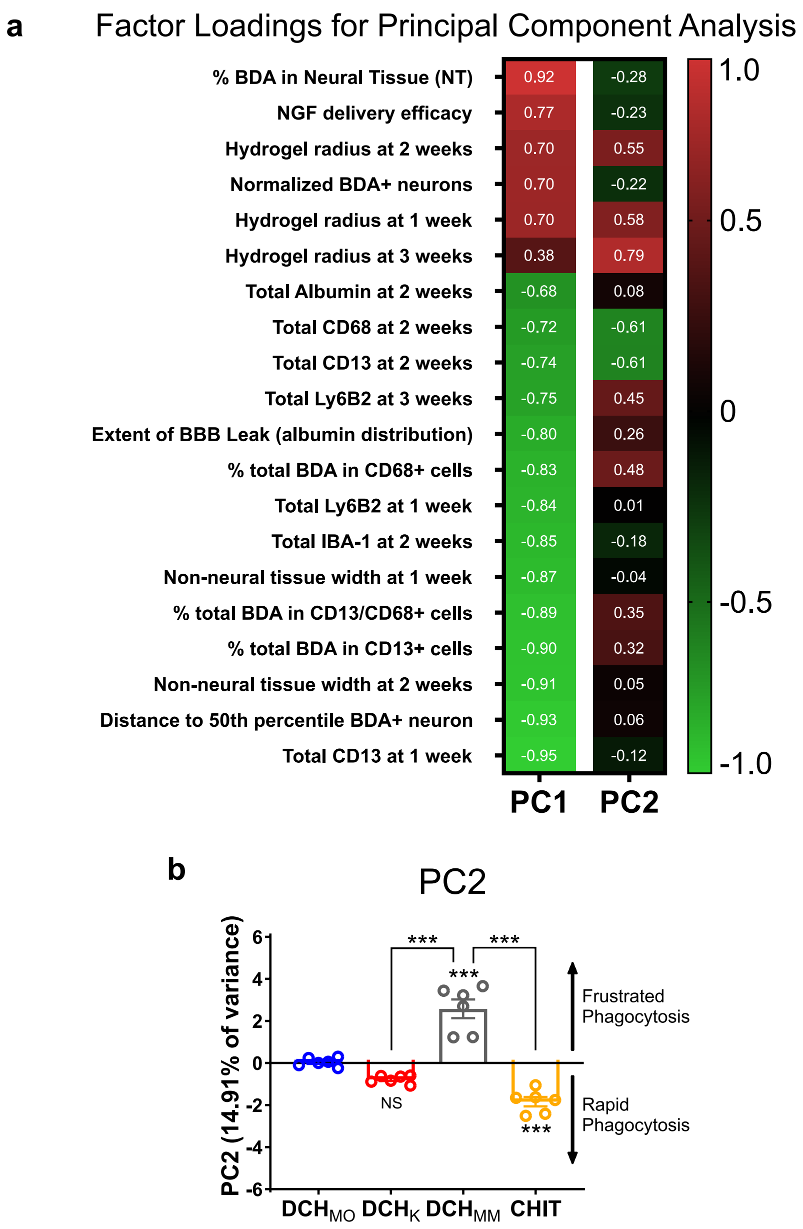


**Supplementary Figure 19 | Principal Component Analysis (PCA) to assess hydrogel function in the CNS. a**. List of parameters used in the PCA with their corresponding factor loadings for the orthogonal PC1 and PC2 axes. The parameters included in the PCA had a Kaiser-Meyer-Olkin measure of sampling adequacy (KMO) of greater than 0.5. **b.** Graphical representation of the positions of the different hydrogel samples along the PC2 axis which accounts for 14.91% of the total variance. NS for $DCH_K$, ***$P < 0.0003$ for $DCH_{MM}$ or CHIT versus $DCH_{MO}$ and ***$P < 0.0001$ for $DCH_K$ or CHIT versus $DCH_{MM}$. Data are mean ± s.e.m with individual data points showing *n* mice per group.
